# An *Arabidopsis* SR protein relieving ABA inhibition of seedling establishment represses ABA-responsive alternative splicing

**DOI:** 10.1101/2023.12.19.572415

**Authors:** Tom Laloum, Guiomar Martín, Martin Lewinski, Romana J. R. Yanez, Tino Köster, Dorothee Staiger, Paula Duque

**Affiliations:** Instituto Gulbenkian de Ciência, Oeiras, Portugal; Department of Biology, Healthcare and the Environment, Faculty of Pharmacy and Food Sciences, University of Barcelona, Barcelona, Spain; RNA Biology and Molecular Physiology, Faculty of Biology, Bielefeld University, Bielefeld, Germany

## Abstract

The phytohormone abscisic acid (ABA) inhibits postgerminative growth under unfavorable conditions to delay the transition to the autotrophic stage and promote plant survival. While stress-induced ABA accumulation is well established to trigger extensive transcriptional changes, it is becoming clear that it also relies on alternative splicing to enhance stress tolerance. However, the upstream components modulating posttranscriptional regulation of the ABA response remain largely unknown. Here, we show that loss of function of the *Arabidopsis* SR34a protein enhances sensitivity to ABA during seedling establishment. Individual-nucleotide crosslinking and immunoprecipitation (iCLIP) combined with RNA-sequencing revealed that SR34a is an alternative splicing regulator that binds predominantly GCU-rich exonic sequences near splice sites. We find that SR34a targets all alternative splicing event types, including in RNAs encoding known determinants of ABA sensitivity, to prevent ABA-responsive splicing in germinated seeds. Our study sheds mechanistic light on how plant SR proteins regulate alternative splicing and counteract ABA inhibition of early plant growth.

## Introduction

The abscisic acid (ABA) hormone is a critical component of plant responses to pervasive environmental stresses such as drought and high salinity. It accumulates in plant cells under osmotic stress, triggering downstream signaling to implement a range of physiological responses to enhance stress tolerance^1^. During early seedling development, ABA plays a central role in inhibiting growth under unfavorable conditions, namely by halting development after seed germination and prior to cotyledon emergence^2^. This postgermination growth arrest is a vital checkpoint restricting the transition from heterotrophic growth to the development of more stress-susceptible autotrophic tissues and is therefore crucial to promote plant survival under stress. Transcriptional, and more recently epigenetic^3^, regulation of this early growth arrest, namely via modulators of seed ABA responses such as ABI5^2^ and ABI3^4^, has been reported but its posttranscriptional control remains elusive despite the fact that alternative splicing has emerged as a key regulatory layer of ABA-mediated stress responses^5^.

RNA splicing is carried out by the spliceosome, a large molecular complex that recognizes splice sites at the intron-exon boundaries^6^. Alternative splicing occurs when splice sites are differentially recognized and multiple transcripts are produced from the same gene. This can result in posttranscriptional regulation of gene expression levels when the alternative splicing event occurs in untranslated (UTR) regions^7^ or lead to the production of alternative protein isoforms if the event is in the coding sequence (CDS). Alternatively-spliced sequences in the CDS can also introduce premature termination codons (PTCs), thereby generating truncated proteins or triggering degradation of the mRNA through nonsense-mediated decay (NMD)^8^.

The selection of splice sites, and hence alternative splicing, is largely influenced by accessory spliceosomal RNA-binding proteins. Amongst them, serine/arginine-rich (SR) proteins comprise a highly conserved family of splicing regulators containing one or two N-terminal RNA recognition motifs (RRMs) and a C-terminal RS domain rich in arginine and serine dipeptides. The molecular function of SR proteins has been well established in animal systems, where they act as major modulators of splice site selection through RRM-dependent binding of regulatory sequences in the pre-mRNA and recruitment of core spliceosome components to nearby splice sites via protein interactions with the RS domain^9^. Apart from splicing, animal SR proteins have also been shown to influence other steps of gene expression, including transcription and mRNA polyadenylation, export, stability, translation or NMD^9^.

The *Arabidopsis thaliana* genome encodes 18 SR proteins classified in six subfamilies. Members of the SR, RSZ and SC subfamilies are orthologs of the human SRSF1, SRSF2 and SRSF7 proteins, respectively, while the SCL, RS2Z and RS subfamilies are plant-specific^10^. Additionally, two *Arabidopsis* SR-like proteins, SR45 and SR45a, display a noncanonical protein structure, with a single RRM flanked by two RS domains^10^. Several *Arabidopsis* SR and SR-like proteins interact with core spliceosomal components, including U1-70K and U2AF^11–16^, and profiling of the transcriptomic defects imposed by SR mutations in plants has substantiated a role for these RNA-binding proteins in splicing regulation^14,17–19^. Moreover, transcripts associated with the *Arabidopsis* SR45 protein were identified by RNA-immunoprecipitation sequencing (RIP-seq) and putative binding motifs inferred from sequence enrichment^17,20^, but mapping of the exact sequences recognized by plant SR proteins using available cutting-edge techniques such as individual-nucleotide resolution crosslinking and immunoprecipitation (iCLIP) is yet to be reported. A molecular function in splicing has been demonstrated for several *Arabidopsis* SR and SR-like proteins. Indeed, RSZ22 binds *in vitro* to RNA sequences known to be recognized by mammalian SR proteins and efficiently complements splicing deficiencies of SRSF7-depleted cell extracts^21^. Moreover, the SCL33 protein binds to the third intron of its pre-mRNA to autoregulate alternative splicing^22^, and SR45 associates with thousands of transcripts in seedlings^20^ and inflorescences^17^, some of which show splicing defects in an *SR45* loss-of-function mutant.

Consistent with a role for alternative splicing in plant stress tolerance, functional links between *Arabidopsis* SR proteins and the response to ABA and related stresses have emerged. Mutations in the *RS40* and *RS41* genes cause hypersensitivity to ABA and high salinity during seed germination and root elongation^23^, while the SCL30a protein represses seed ABA sensitivity during germination under salt stress^24^ and SR45a negatively regulates seedling tolerance to salt^18^. The well characterized SR45 binds transcripts enriched in ABA signaling functions^20^, represses glucose-induced ABA accumulation and signaling^25,26^, and its loss-of-function results in altered seedling sensitivity to the hormone^25–27^. It is also noteworthy that RSZ22 appears to be a dephosphorylation target of the *Arabidopsis* phosphatase 2C HIGHLY ABA-INDUCED 1 (HAI1), a key component of the ABA signaling^28^. While these data disclose a role in modulating the ABA pathway, the RNA molecules and sequences targeted by SR proteins to regulate stress responses mediated by the hormone are unknown.

SR34a is one of the four *Arabidopsis* orthologs of the human SRSF1 splicing factor^10^ and, apart from a previous report that it exhibits nucleocytoplasmic shuttling^29^, remains uncharacterized. Here, we show that loss of SR34a function enhances plant sensitivity to ABA and osmotic stress during the developmental transition to autotrophic growth. We successfully implemented iCLIP2^30,31^ in *Arabidopsis* germinated seeds, and combined it with RNA-sequencing (RNA-seq) profiling of a loss-of-function mutant to identify the transcripts directly regulated by SR34a. This revealed a major function for SR34a as a regulator of alternative splicing that binds its target RNAs by recognizing predominantly exonic GCU sequences in the vicinity of splice sites. Notably, SR34a directly represses most of the ABA-induced alternative splicing changes during seedling establishment, including in known regulators of the ABA response. Our results shed new light on SR protein modulation of plant alternative splicing, underlining its impact on ABA-mediated control of postgerminative growth.

## Results

### The SR34a protein negatively regulates ABA sensitivity during early seedling development

Publicly available data (e.g. https://bar.utoronto.ca/eplant/, https://pastdb.crg.eu/) indicated that expression of the *Arabidopsis* SR protein gene *SR34a* (AT3G49430) is induced during seed germination. Our analysis of the *SR34a* expression pattern using RT-qPCR showed that transcript levels were significantly enhanced during seed imbibition at 4°C (stratification) and confirmed a sharp increase early in the germination process, with expression peaking at 24 hours and decreasing thereafter (Fig. 1a).

**Fig. 1.**
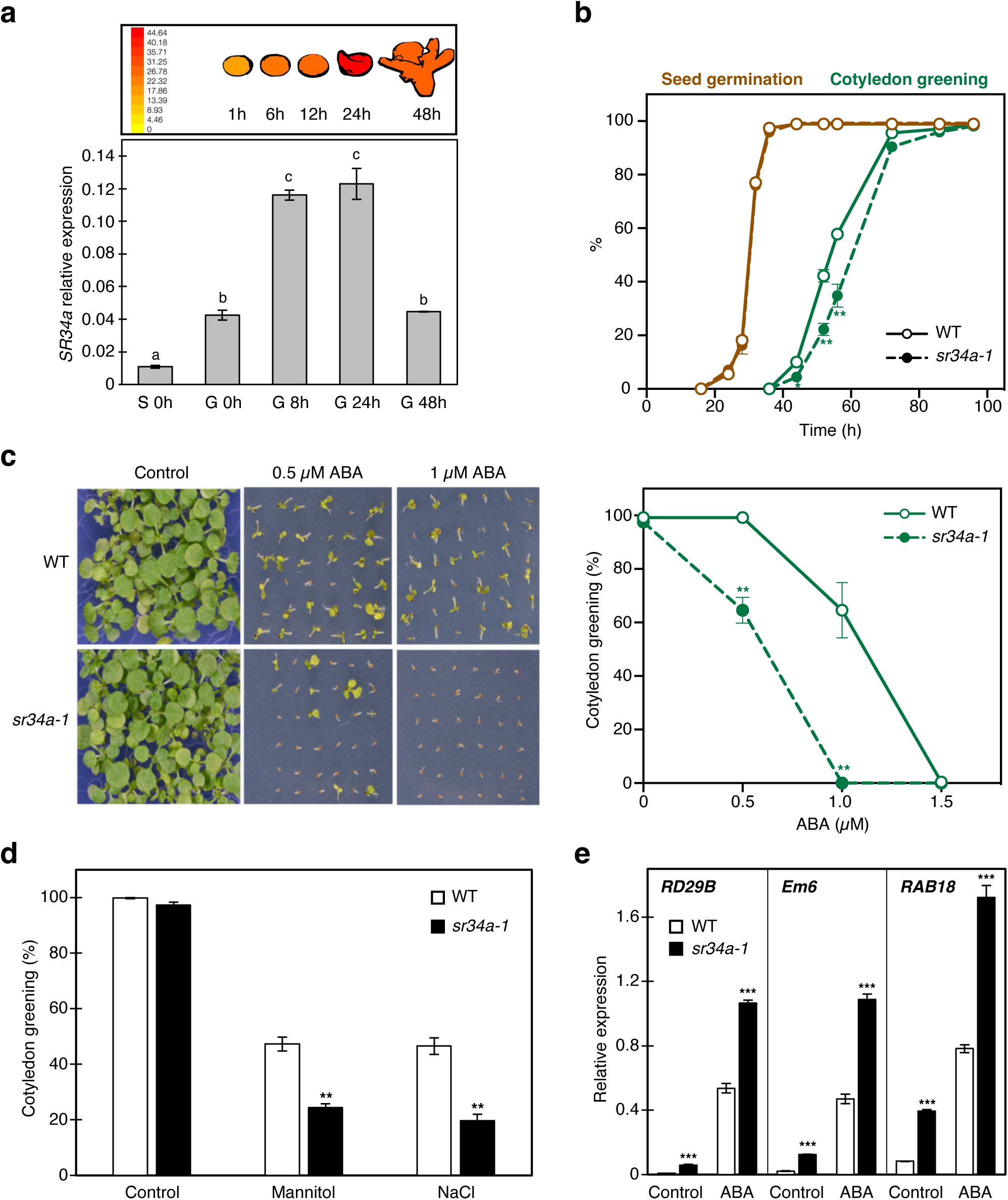
ABA phenotypes of the *sr34a-1* mutant during early seedling development. **a** RT-qPCR analysis of *SR34a* transcript levels at the onset of seed stratification (S 0h) and at 0, 8, 24 or 48 h of the germination process (G 0h, G 8h, G 24h, G 48h). Letters indicate statistically significant differences between timepoints (p < 0.05; Two-way ANOVA with Tukey HSD test). The top panel is from the ePlant tool (https://bar.utoronto.ca/eplant) and illustrates RNA-seq data on *SR34a* expression in germinating seeds 1, 6, 12, 24 and 48 h poststratification^70^. **b** Seed germination (brown) and cotyledon greening (green) percentages of Col-0 wild-type (WT) and *sr34a-1* mutant plants scored during the first 4 d after stratification (means ± SE, *n* = 3). Results are representative of 5 independent experiments. **c** Representative images and quantification of cotyledon greening percentages of Col-0 WT and *sr34a-1* mutant seedlings scored 10 d after stratification and growth under different ABA concentrations (means ± SE, *n* = 3). Results are representative of 9 independent experiments. **d** Cotyledon greening percentages of Col-0 WT and *sr34a-1* mutant seedlings scored 5 d after stratification and growth in medium supplemented or not with 300 mM mannitol or 170 mM NaCl (means ± SE, *n* = 3). Results are representative of 3 independent experiments. **e** RT-qPCR analysis of *RD29B*, *Em6* and *RAB18* transcript levels in Col-0 WT and *sr34a-1* mutant seeds germinated for 44 h and submitted to a 2-h mock (control) or 0.5 µM ABA treatment (means ± SE, *n* = 4). In **b-e**, asterisks indicate statistically significant differences from the wild type under each condition or at each timepoint (* p < 0.05, ** p < 0.01, *** p < 0.001; Student’s *t*-test).

To uncover the biological roles of SR34a, we isolated a homozygous T-DNA mutant line, *sr34a-1*, carrying the insertion in the gene’s twelfth exon (Supplementary Fig. 1a). RT-qPCR using primers flanking the T-DNA insertion revealed near depletion of the full-length *SR34a* transcript, while primers annealing upstream of the insertion pointed to the production of a truncated transcript whose expression is only ∼25% of the full-length transcript in the wild-type (Supplementary Fig. 1b). The *sr34a-1* mutation is thus expected to severely affect SR34a function.

Given the marked induction of *SR34a* expression during seed germination, we first asked whether the *sr34a-1* mutant would be affected in the earliest steps of plant development. As seen in Figure 1b, while mutant seeds germinated as fast as the wild type, a slight but significant delay was detected during greening of the cotyledons. As early plant development is tightly controlled by ABA and other SR and SR-like proteins have been implicated in ABA-mediated processes^18,23–25^, we next assessed the *sr34a-1* mutant’s response to exogenous ABA. Figure 1c shows that the mutant displayed strong hypersensitivity to the hormone during postgerminative growth, with ABA concentrations that allowed 60% of wild-type seedlings to develop green cotyledons arresting growth of the mutant upon seed germination. In agreement with the established role of ABA in mediating plant responses to osmotic stress, greening of *sr34a-1* cotyledons was significantly delayed under elevated concentrations of mannitol (which mimics drought stress) or NaCl (Fig. 1d).

To check that the mutant phenotype was caused by loss of SR34a function, two independent transformant lines, C1 and C2, expressing a GFP-tagged version of SR34a driven by its endogenous promoter in the mutant background were isolated. After confirming that the transgene was expressed at wild-type levels (Supplementary Fig. 2a), we found that both transgenic lines displayed full rescue of the mutant’s ABA hypersensitivity (Supplementary Fig. 2b). Moreover, microscopy analyses confirmed nuclear localization of SR34a, with no evident change upon ABA treatment (Supplementary Fig. 2c).

Finally, to confirm ABA hypersensitivity of the *sr34a-1* mutant at the molecular level, we analyzed the expression of the ABA-responsive genes *RD29B*, *Em6* and *RAB18* in wild-type and *sr34a-1* germinated seeds upon a transient mock or ABA treatment. In line with the enhanced *sr34a-1* response during ABA-mediated processes, the expression levels of all three marker genes were significantly higher in the mutant both under control and ABA conditions (Fig. 1e).

Thus, the *Arabidopsis SR34a* gene is markedly induced during seed imbibition and germination. Reverse genetics and molecular analyses revealed that the encoded SR protein is a negative regulator of ABA-mediated developmental, stress and transcriptional responses during early seedling development.

### SR34a downregulates ABA induction of gene expression

The higher expression of three ABA marker genes in the *sr34a-1* mutant prompted us to investigate the global transcriptomic effects of the SR34a protein during postgerminative growth and in the ABA response. To this end, we conducted an RNA-seq experiment in 44-hour germinated Col-0 wild-type and *sr34a-1* mutant seeds submitted to a 2-h mock (control) or ABA treatment. The effectiveness of the latter was confirmed by gene ontology (GO) analysis of the 587 genes whose expression was altered by ABA in the wild type (fold change ≥ 2; Fig. 2a; Supplementary Data 1), which showed strong enrichment in functional categories such as response to ABA or water deprivation (Supplementary Data 2). In the *sr34a-1* mutant, ABA regulated 654 genes, 517 of which were also regulated in the wild type (Fig. 2a; Supplementary Data 1), revealing that the SR34a protein hardly affects the set of genes that responds to ABA in germinated *Arabidopsis* seeds.

**Fig. 2.**
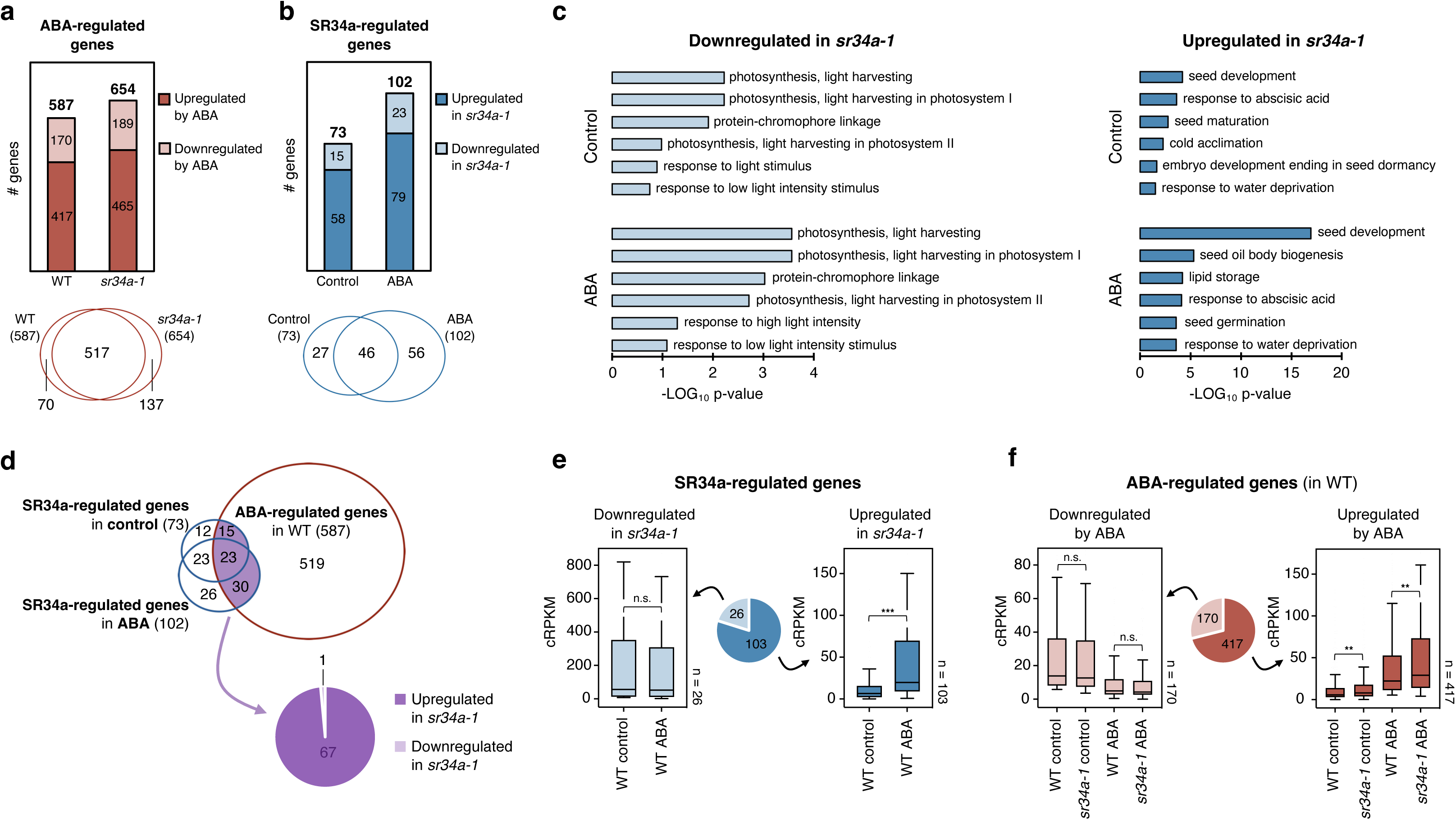
Gene expression changes associated with loss of SR34a function and ABA exposure. **a** Number of genes up-(dark red) or down-(light red) regulated by ABA and overlap between the genes regulated by ABA in the wild type (WT) and *sr34a-1*. **b** Number of genes up-(dark blue) or down-(light blue) regulated in the *sr34a-1* mutant and overlap between the genes regulated by SR34a under control and ABA conditions. **c** Six most enriched Gene Ontology (GO) biological process categories for the genes up- or downregulated in the *sr34a-1* mutant under control or ABA conditions. **d** Overlap between ABA-regulated genes (in the WT) and SR34a-regulated genes under control or ABA conditions, and number of genes upregulated or downregulated among those altered both in *sr34a-1* and by ABA (in the WT). **e** Expression levels of the genes up-(dark blue) or down-(light blue) regulated in the *sr34a-1* mutant in WT samples under control or ABA conditions. **f** Expression levels of the genes up-(dark red) or down-(light red) regulated by ABA in WT or *sr34a-1* samples under control or ABA conditions (right). In **e** and **f**, asterisks indicate statistically significant differences (** p < 0.01, *** p < 0.001; Mann & Whitney test). n.s., not significant.

We next focused on comparing the wild-type and *sr34a-1* transcriptomes and identified 73 or 102 differentially-expressed genes (DEGs, fold change ≥ 2) in the mutant upon control or ABA treatments, respectively, finding a 46-gene overlap between the two sets of DEGs (Fig. 2b; Supplementary Data 3). Thus, SR34a is already largely active under basal conditions and modulates gene expression under both basal and ABA conditions. Consistent with the observed delay in cotyledon greening (see Fig. 1b, c), categories related to photosynthetic processes were strongly enriched among the genes downregulated in the *sr34a-1* mutant, many of which encoded chlorophyll-binding proteins (Fig. 2c; Supplementary Data 3, 4). By contrast, genes upregulated in *sr34a-1* were strongly enriched in functions related to seed development and maturation as well as to the response to ABA or water deprivation (Fig. 2c; Supplementary Data 3, 4).

To understand SR34a regulation of ABA-responsive gene expression, we assessed the overlap between the genes regulated by SR34a and those responding to ABA in wild-type germinated seeds. As shown in Figure 2d, over half of the genes that were changed in the *sr34a-1* mutant under control (38/73) or ABA (53/102) conditions were also changed by the ABA treatment in the wild type, indicating that SR34a modulates the expression of ABA-responsive genes under both basal and ABA conditions. Furthermore, 67 of the 68 genes whose expression was affected by both the *sr34a-1* mutation and ABA were upregulated in the mutant (Fig. 2d), indicating that unlike downregulated genes those that are upregulated in the *sr34a-1* mutant are predominantly ABA responsive.

To gain deeper insight into the reciprocal regulation of gene expression by SR34a and ABA, we asked how the phytohormone modulates the expression of SR34a-regulated genes and, conversely, how the SR protein regulates ABA-responsive genes. Unlike genes downregulated in *sr34a-1*, whose expression was not affected by ABA, genes upregulated in the mutant were expressed at a significantly higher level in the ABA-treated wild type when compared with control conditions (Fig. 2e; Supplementary Fig. 3). Reciprocally, ABA-induced genes were expressed at a significantly higher level in *sr34a-1*, while ABA-repressed genes were not differentially expressed in the mutant (Fig. 2f). These results indicate that specifically ABA-induced genes exhibit enhanced expression in the *sr34a-1* mutant.

Therefore, although SR34a does not appreciably affect the identity of the genes responding to ABA in *Arabidopsis* germinated seeds, it represses the expression of genes induced by the hormone under both control and ABA conditions, in clear agreement with its negative regulation of ABA sensitivity during early seedling development.

### SR34a regulates alternative splicing similarly under basal and ABA conditions

Given that SR proteins are key regulators of alternative splicing, we next asked whether loss of *SR34a* function would cause significant defects in this process. Comparison of wild-type and *sr34a-1* RNA-seq samples identified a total of 242 differential alternative splicing (DAS) events, i.e. showing a difference in Percent Spliced In (ΔPSI) > 15, with 137 and 135 events detected under control and ABA conditions, respectively, and 30 DAS events being common to the two conditions (Fig. 3a; Supplementary Data 5). The alternative splicing event types affected by the mutation were similar in control and ABA conditions, the vast majority being intron retention (IR, ∼35%) and alternative 3’ splice site (Alt3, ∼35%), followed by exon skipping (ES, ∼20%) and alternative 5’ splice site (Alt5, ∼10%) (Fig. 3a). Remarkably, only one of the 242 events occurred in a gene (AT4G28680) whose expression also differed between the wild type and the mutant (Supplementary Data 3, 5).

**Fig. 3.**
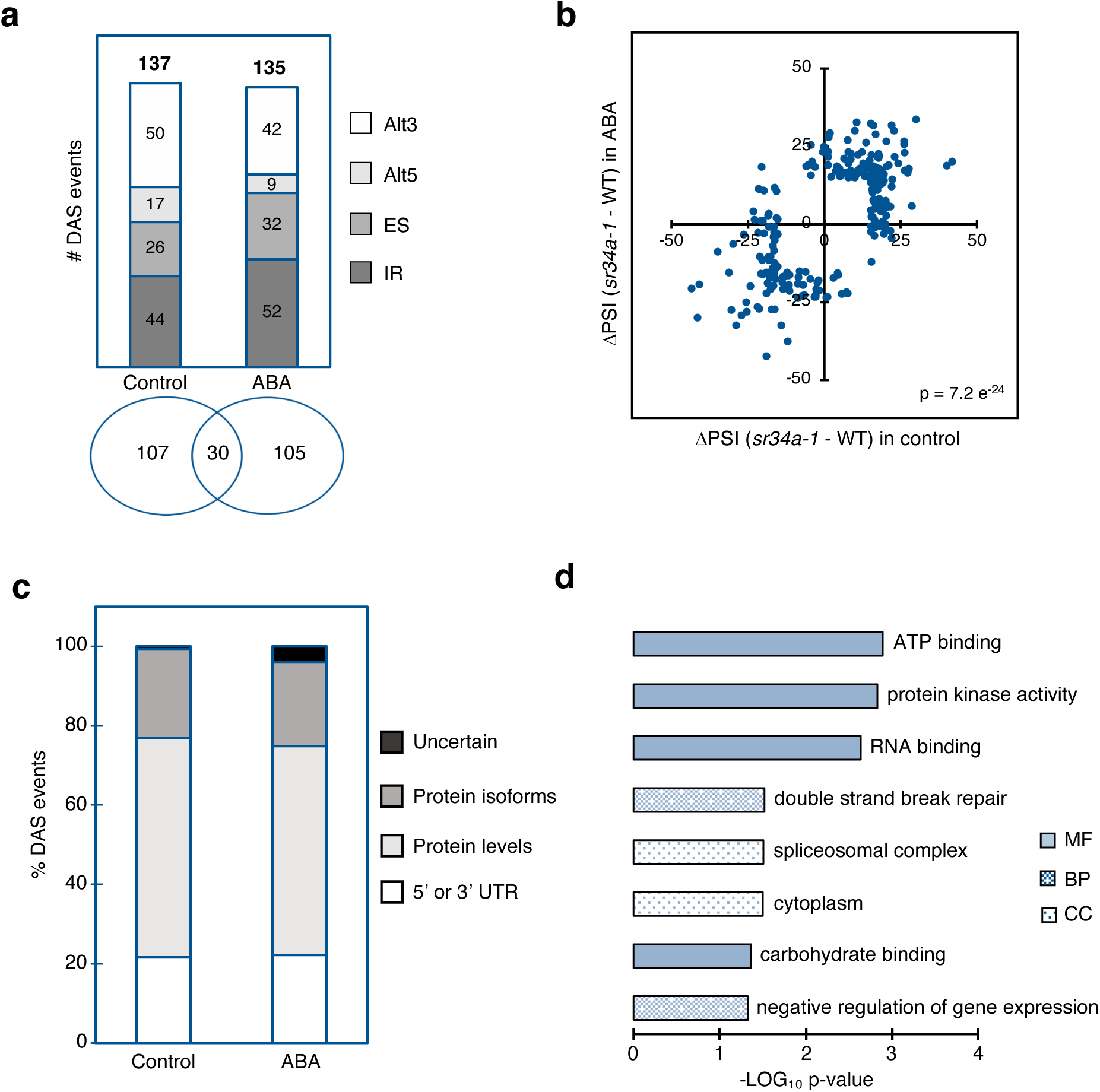
Alternative splicing changes associated with loss of SR34a function. **a** Total number of differential alternative splicing (DAS) events in the *sr34a-1* mutant under control and ABA conditions, including numbers for each of the four major types of alternative splicing events, and overlap between the DAS events identified under control and ABA conditions. IR, intron retention; ES, exon skipping; Alt5, alternative 5’ splice site; Alt3, alternative 3’ splice site. **b** Effect (differential inclusion levels) of the *sr34a-1* mutation on the 242 DAS events in control or ABA conditions. The p-value indicates statistical significance of the association between the ΔPSI in control and ABA conditions (Fisher’s exact test with a 2 x 2 contingency table). **c** Percentage of DAS events occurring in UTRs or in the protein-coding sequence. Among the latter, the percentage of events that potentially affect protein levels (by including or excluding a premature stop codon) or produce alternative protein isoforms is indicated. UTR, untranslated region. **d** All significantly-enriched (p < 0.05) Gene Ontology (GO) categories for the 197 genes displaying DAS events under control and/or ABA conditions. MF, molecular function; BP, biological process; CC, cellular compartment.

To assess the effect of ABA on the splicing function of SR34a, we quantified the alternative splicing changes imposed by the mutation (ΔPSI*_sr34a-1_* _-wild type_) under each condition. Despite the relatively low overlap between control and ABA conditions (Fig. 3a), we found that globally the DAS events were similarly regulated by SR34a, i.e. alternative sequences more included in the mutant under control conditions were also more included upon ABA exposure, and conversely sequences less included in *sr34a-1* in the absence of ABA were also less included after hormone treatment (p < 0.0001; Fig. 3b). This indicates that ABA does not substantially alter regulation of splicing by SR34a.

Next, to gain insight into the functional relevance of SR34a-regulated alternative splicing, we assessed the predicted impact of the DAS events on the mature transcript (see Methods for details). Under both control and ABA conditions, ∼20% of the DAS events occurred in UTR regions and may have regulatory consequences, e.g. by affecting transcript stability or translation efficiency (Fig 3c). The majority of the remaining events taking place in the coding region are likely to change the proportion of unproductive transcripts (by including or excluding a premature termination codon, leading to nonsense-mediated RNA decay) rather than generate different protein isoforms (Fig. 3c).

The alternative splicing changes detected by RNA-seq were validated via semi-quantitative RT-PCR for five randomly-selected DAS events exhibiting a substantial change in the *sr34a-1* mutant (|ΔPSI*_sr34a-1_*_-wild type_| > 20) under at least one condition. As shown in Supplementary Figure 4, for all five events, quantification of the PCR products amplified using primers flanking the alternatively-spliced sequence yielded PSI values closely matching those obtained in the RNA-seq experiment.

The 242 SR34a-regulated alternative splicing events occurred in a total of 196 genes. Notably, these genes were strongly enriched for ATP-binding functions (Fig. 3d, Supplementary Data 6), which included mostly kinases, but also proteins with very diverse biological roles such as ATPases, ABC transporters, proteins involved in DNA repair or in lipid biosynthesis. Many of the DAS events also occurred in transcripts encoding RNA-binding proteins involved in splicing regulation, RNA 3’ end processing, RNA degradation or in the reading of m6A marks. In particular, the DAS events affected several SR and SR-like genes, including *SR30* (AT1G09140), *SR34b* (AT4G02430), *SR45a* (AT1G07350) or *RS31A* (AT2G46610), as well as *AFC1* (AT3G53570) and *PRP4K* (AT1G13350) coding for protein kinases that regulate SR protein activity^32^, supporting the existence of a splicing regulatory loop controlled by SR34a, as already proposed for several other splicing factors^22,33–35^.

The above results uncover an important role for the SR34a protein in the regulation of alternative splicing during early seedling development. Under both optimal growth conditions and in response to ABA, SR34a controls all types of alternative splicing events in nearly 200 genes encoding mainly proteins with kinase, ATP- and RNA-binding activity.

### SR34a binds GCU-rich exonic sequences near splice sites in vivo

Identifying the transcripts and sequence motifs targeted in vivo by RNA-binding proteins is essential to understand their mode of action. Crosslinking and immunoprecipitation (CLIP) analysis is currently the most reliable technique for faithfully deciphering RNA-protein interactions^31,36,37^, but it remains challenging to implement in plant systems. To covalently crosslink RNA targets to the SR34a protein in germinated seeds, we adapted UV-crosslinking protocols established for mammalian cells, *Caenorhabditis elegans* and *Arabidopsis* seedlings^31,36,38^ (see Methods and Fig. 4a). The SR34a-GFP transgenic line (C1; see Supplementary Fig. 2) and control lines expressing the GFP protein alone under the control of the 35S promoter grown under the exact same conditions as for the RNA-seq experiment were used.

**Fig. 4.**
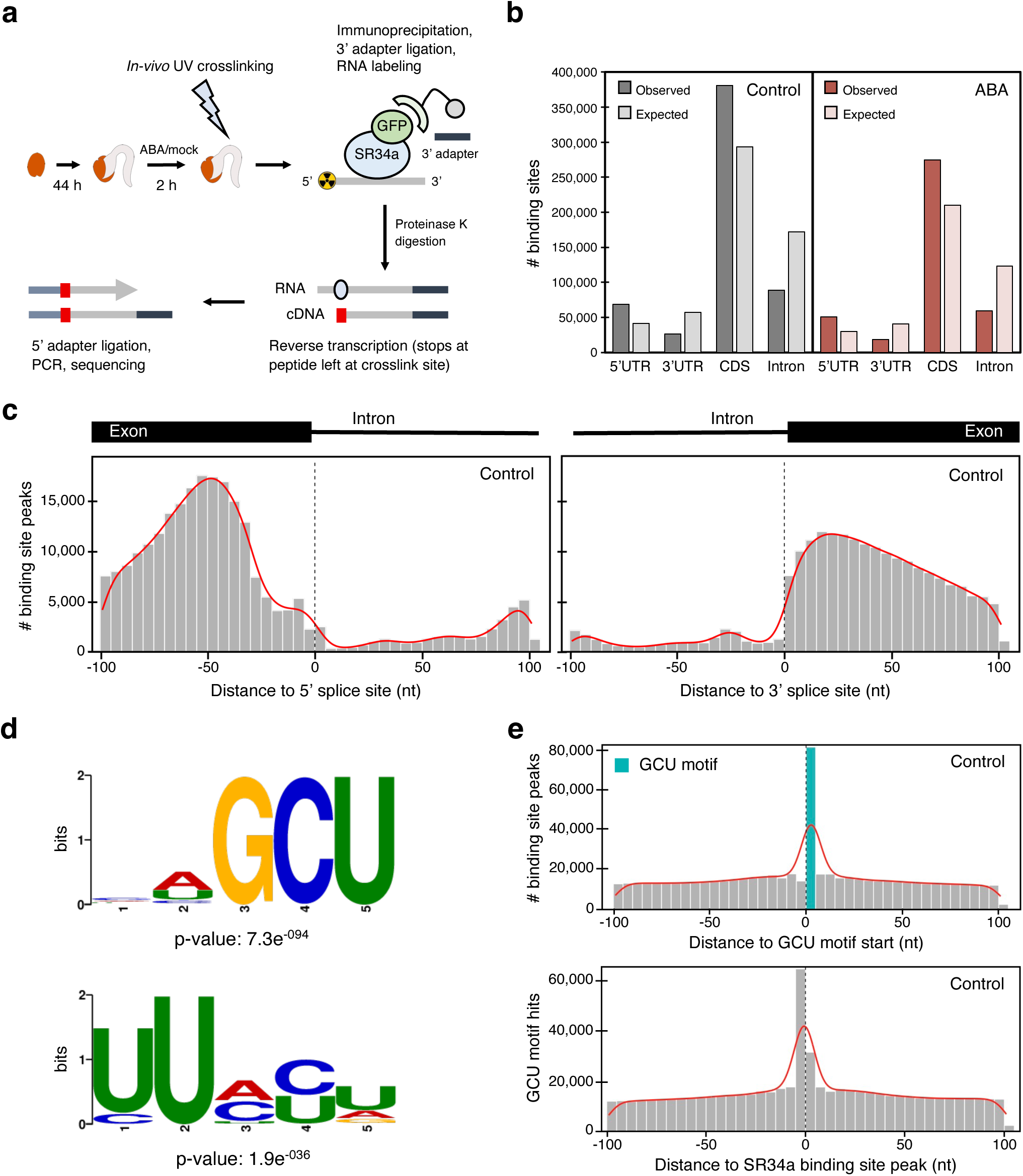
SR34a binding landscape. **a** Diagram illustrating the iCLIP2 protocol adapted to *Arabidopsis* germinated seeds (see Methods for details). **b** Number of reproducible 9-nt binding sites identified in different transcript regions (observed) versus the binding site distribution predicted from the cumulative length of the indicated region in the genome based on TAIR10 (expected) under control (gray) or ABA (red) conditions. **c** Number of SR34a iCLIP binding site peaks (i.e. middle position of binding sites) mapped in the vicinity of 5’ and 3’ splice sites under control conditions. **d** Binding motifs significantly enriched within SR34a binding sites identified under control conditions. **e** Number of iCLIP binding site peaks identified under control conditions as a function of their distance to GCU motifs in the *A. thaliana* genome (top), and number of GCU motif hits as a function of their distance to binding site peaks (bottom).

After confirming an efficient immunoprecipitation of the SR34a-GFP and GFP proteins from lysates of the C1 line and control transgenic plants expressing the GFP alone, respectively (Supplementary Fig. 5a), we monitored the formation of specific RNA-SR34a-GFP complexes. As seen in Supplementary Figure 5b, these were detected in UV-crosslinked germinated C1 seeds under both control and ABA conditions but not in non-crosslinked tissue, demonstrating the suitability of the method to retrieve RNA-protein complexes from *Arabidopsis* seeds as achieved previously with seedlings^31,37,39,40^. Subsequent RNase I treatment eliminated most of the crosslinked RNA, leaving an expected band ∼5 kDa above the protein’s molecular weight (Supplementary Fig. 5b), thus confirming that the RNA signal was specific to the SR34a-GFP protein^38^. After iCLIP2 library preparation and sequencing of the crosslinked RNA from SR34a-GFP and GFP samples (Supplementary Fig. 6), followed by read processing and peak calling at the crosslink sites, reproducible 9-nt SR34a binding sites were identified by extending the called peaks by 4 nt in both directions (see Methods for details). In line with the similar SR34a regulation of alternative splicing in control and ABA conditions (see Fig. 3b), we found a high level of overlap (78%; data not shown) between the binding sites detected in the two conditions, indicating that ABA has a minor impact on the location of SR34a RNA binding. To determine the transcript regions bound preferentially by SR34a, the observed binding site distribution was compared to the distribution expected from the cumulative length of the indicated region in the genome. Figure 4b shows that binding was found to be enriched within the CDS and 5’ UTR regions, while depleted from intronic and 3’ UTR regions.

Studies in animal systems have revealed that SR proteins bind pre-mRNAs primarily within exons, recruiting spliceosomal components to adjacent 5’ and 3’ splice sites^9^. To determine the position of SR34a binding, we monitored the density of iCLIP binding sites at the exon-intron and intron-exon boundaries. Notably, both under control and ABA conditions, SR34a showed a strong preference for binding exonic regions within 100 nt of splice sites, with the highest binding site density observed around 50 nt upstream of the 5’ splice site and 25 nt downstream of the 3’ splice site (Fig. 4c; Supplementary Fig. 7a). As also found previously for mammalian SR proteins^41^, SR34a binding sites were more prevalent upstream of 5’ splice sites than downstream of 3’ splice sites (Fig. 4c; Supplementary Fig. 7a).

Finally, to identify the sequences directly bound by SR34a, we searched for enriched motifs within the binding sites using Sensitive Thorough Rapid Enriched Motif Elicitation (STREME). This revealed two major motifs enriched under both control and ABA conditions (Fig. 4d; Supplementary Fig. 7b) — while detection of U-rich motifs could be due to UV-crosslinking bias towards these sequences^42^, we found an even higher significant enrichment in GCU-containing motifs. Assessment of the number of iCLIP binding site peaks as a function of their distance to GCU sequences or, conversely, the number of GCU motifs as a function of the distance to binding site peaks (Fig. 4e; Supplementary Fig. 7c) confirmed the strong association with GCU motifs.

Our iCLIP analysis thus revealed that the SR34a RNA-binding protein recognizes predominantly GCU motifs, binding at exonic *cis* elements in the vicinity of splice sites as established for animal SR proteins. These binding features corroborate a prevalent role in alternative splicing regulation and appear to be unaffected by the ABA hormone.

### SR34a directly controls alternative splicing

To evaluate whether SR34a directly regulates alternative splicing during early seedling development, we first checked the proportion of SR34a-regulated DAS events identified by RNA-seq that corresponded to transcripts found to be bound by SR34a in the iCLIP analysis. As seen in Figure 5a, these direct DAS (dDAS) events represented the vast majority of DAS events, 74% and 68% under control and ABA conditions, respectively (see also Supplementary Data 7, 8), substantiating a major role for SR34a as a splicing factor.

**Fig. 5.**
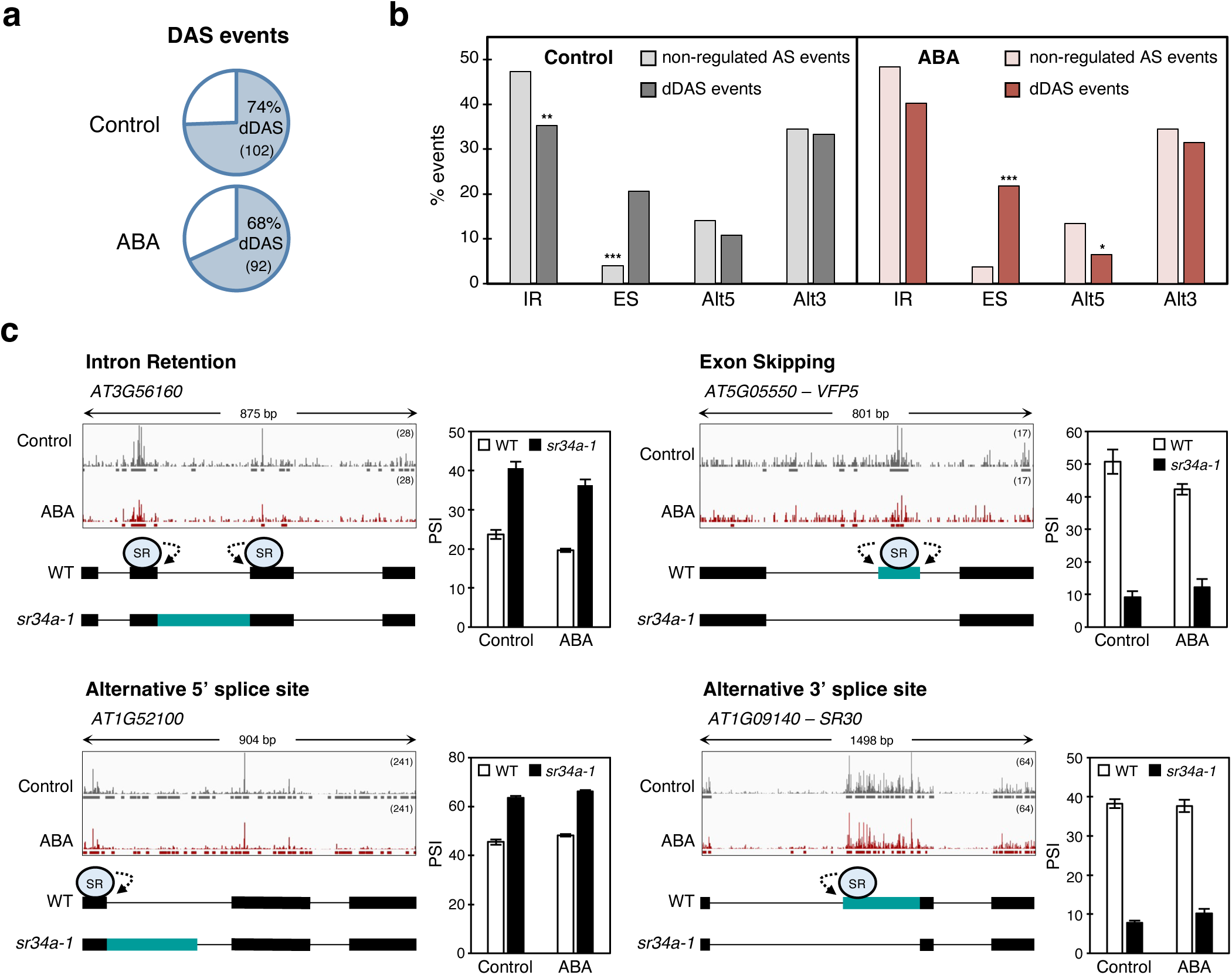
Direct SR34a splicing targets. **a** Percentage of differential alternative splicing (DAS) events in the *sr34a-1* mutant whose transcripts are bound by SR34a (dDAS). The number of events for each category is shown in parentheses. **b** Percentages of intron retention (IR), exon skipping (ES), alternative 5’ splice site (Alt5) and alternative 3’ splice site (Alt3) among the 102 or 92 dDAS events identified under control (dark gray) or ABA (dark red) conditions, respectively. The lighter colored bars (non-regulated AS events) indicate the percentages of each type of alternative splicing event among all the events detected in the transcriptome that were unaffected by the *sr34a-1* mutation (see Supplementary Data 9, 10). Asterisks indicate significant differences (* p < 0.05, ** p < 0.01, *** p < 0.001; hypergeometric test). **c** SR34a iCLIP peaks and RNA-seq PSI values for one example of each type of dDAS event. The number of crosslink sites is indicated in parentheses, and significant binding sites identified under control or ABA conditions are represented respectively by gray and red horizontal lines under the iCLIP peaks. Model diagrams for SR34a control of the dDAS events are also shown, with the alternatively-spliced sequence indicated in teal blue and the arrows representing regulation of splice site selection.

To better understand the splicing function of SR34a, we next compared the different types of dDAS events with a set of alternative splicing events, named “non-regulated AS events”, whose inclusion did not change substantially between the wild type and *sr34a-1* (10 < PSI < 90, ΔPSI*_sr34a-1_*_-wild type_ < 5) (Fig. 5b: Supplementary Data 9, 10). Consistent with recent reports of global *A. thaliana* alternative splicing patterns^43,44^, we found that under both control and ABA conditions IR was the most prevalent alternative splicing event type in the “non-regulated AS events” sets, followed by Alt3, Alt5 and ES (Fig. 5b). Among the dDAS events, we detected the largest numbers for IR and Alt3 events, with SR34a regulating directly also ES and, to a lower extent, Alt5 events (Fig. 5b). The majority of IR events (25/36 and 28/37 under control and ABA conditions, respectively) were preferentially changed towards more intron inclusion in the *sr34a-1* mutant (Supplementary Fig. 8), indicating that the SR34a protein promotes splicing of these introns. Under both conditions, ES events were significantly enriched among the dDAS events when compared with the proportion of non SR34a-regulated ES events in the transcriptome (Fig. 5b), pointing to preferential targeting of this type of event by SR34a. Moreover, while under control conditions the SR protein promoted or repressed ES to a similar extent, an intriguingly high proportion of alternative exons was found to be included in the *sr34a-1* mutant upon ABA treatment (Supplementary Fig. 8), suggesting that SR34a tends to inhibit exon inclusion in the presence of the hormone.

We next examined the iCLIP binding sites associated with dDAS transcripts, again detecting general exonic binding at the vicinity of the alternative-spliced sequences, as illustrated by the examples in Figure 5c. For instance, SR34a enhancement of removal of an intron from the *At3g56160* transcript was associated with exonic iCLIP binding sites close to the intron’s 5’ and 3’ splice sites (Fig. 5c, top left), while enhanced inclusion of an exon in *At5g05550* coincided with the presence of SR34a binding sites within the exon (Fig. 5c, top right). For SR34a regulation of Alt5 and Alt3 events in *At1g52100* and *At1g09140*, respectively, splice site selection correlated with exonic binding of the SR protein near the preferred splice site (Fig. 5c, bottom).

In brief, integration of RNA-seq and iCLIP data revealed that SR34a largely binds RNA to regulate alternative splicing, again pointing to a major role in influencing splice site selection by binding near 5’ and 3’ splice sites. While IR is the type of alternative splicing event most frequently regulated by SR34a, the protein appears to disproportionally target ES.

### SR34a represses ABA-inducible alternative splicing changes

Thus far, our analyses of the RNA-seq and iCLIP data revealed details on the mode of action and identified the direct splicing targets of SR34a under both basal and ABA conditions, but provided little molecular insight into its control of ABA sensitivity during early seedling development. To evaluate the effect of SR34a on ABA-induced splicing changes, we turned to comparing control and ABA samples within each genotype. As expected, we found that the hormone changed the splicing pattern of multiple genes also in germinated *Arabidopsis* seeds. We first asked whether the splicing changes induced by ABA (in the wild type) were similar to those caused by the *sr34a-1* mutation during postgerminative growth. As seen in the scatter plot of Figure 6a, alternatively-spliced sequences showing more inclusion in response to ABA in wild-type seeds were in general also more included in *sr34a-1* under control conditions, and conversely sequences less included in response to ABA showed less inclusion in the mutant. In other words, ABA-mediated splicing changes positively correlated with those imposed by the mutation under basal conditions, indicating that the SR34a protein represses ABA-inducible alternative splicing prior to hormone exposure.

**Fig. 6.**
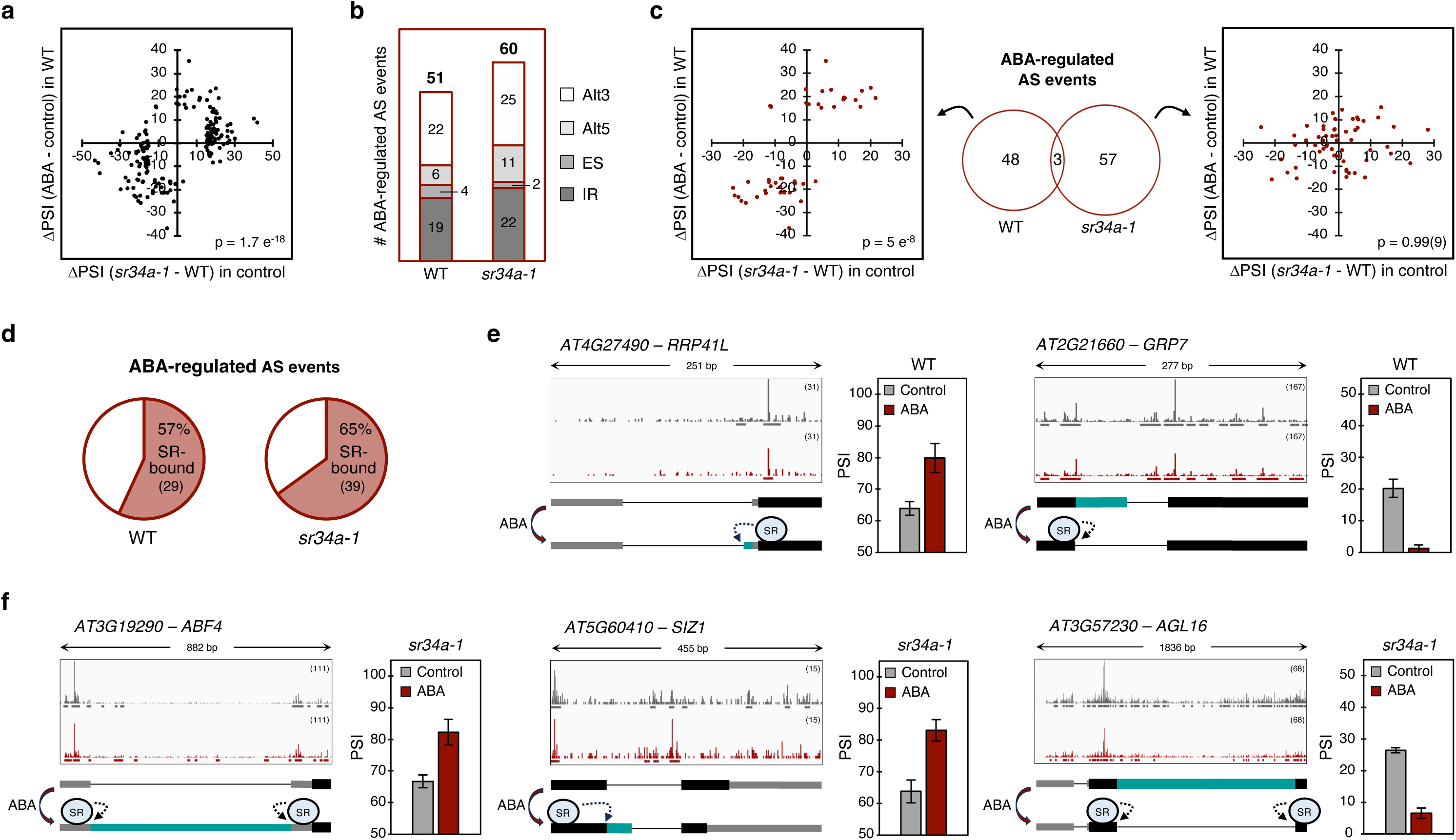
Effect of SR34a on ABA-responsive alternative splicing. **a** Effect (differential inclusion levels) of ABA or the *sr34a-1* mutation on all alternative splicing events found to be either ABA responsive (in the wild type) or SR34a regulated (in basal conditions). **b** Total number of ABA-regulated alternative splicing events in the WT and the *sr34a-1* mutant, including numbers for each of the four major types of alternative splicing event. IR, intron retention; ES, exon skipping; Alt5, alternative 5’ splice site; Alt3, alternative 3’ splice site. **c** Overlap between the ABA-regulated alternative splicing events in the WT and the *sr34a-1* mutant as well as effect (differential inclusion levels) of ABA or the *sr34a-1* mutation on the 48 alternative splicing events responding to ABA only in the WT (left) or on the 57 events changed by ABA only in the *sr34a-1* mutant (right). **d** Percentage of ABA-regulated alternative splicing events in the WT or the *sr34a-1* mutant whose transcripts are bound by SR34a. The number of events for each category is shown in parentheses. **e,f** SR34a iCLIP peaks and RNA-seq PSI values for five known regulators of ABA sensitivity showing ABA-mediated splicing changes only in the WT (**e**) or in *sr34a-1* (**f**) germinated seeds. The number of crosslink sites is shown in parentheses, and significant binding sites identified under control or ABA conditions are represented respectively by gray and red horizontal lines under the iCLIP peaks. Model diagrams for SR34a control of the ABA-regulated alternative splicing events are also shown, with the alternatively-spliced sequence indicated in teal blue and the arrows representing regulation of splice site selection. In **a** and **c**, p-values indicate statistical significance of the association between the depicted variables (Fisher’s exact test with a 2 x 2 contingency table).

Our comparison of control and ABA RNA-seq data identified significant differences for 51 alternative splicing events in the wild type and 60 in the *sr34a-1* mutant (Fig. 6b, Supplementary Data 11). Loss of *SR34a* function does not appear to affect the type of alternative splicing events regulated by ABA, as similar proportions of each event type were found in the two genotypes, with Alt3 (42-43%) and IR (37%) being the most affected by the hormone (Fig. 6b, Supplementary Data 11). Notably, none of the alternative splicing changes induced by ABA in the wild type and only one in *sr34a-1* occurred in a gene (AT1G54100) whose expression was also affected by ABA.

In striking contrast with gene expression, where ABA affected a similar set of genes in wild-type and *sr34a-1* germinated seeds (see Figure 2a), the hormone induced a very different response in the two genotypes at the alternative splicing level, with only three shared events between the 51 and 60 regulated by ABA in the wild type and the mutant, respectively (Fig. 6c Supplementary Data 11). While the 48 alternative splicing events significantly changed in the wild type but not in the mutant are expected to have their ABA regulation mediated by SR34a, we found an even larger set of events significantly changed by ABA only in *sr34a-1* (Fig. 6c), indicating that the SR34a protein represses ABA regulation of the latter alternative splicing events. In line with the global tendency shown by all alternative splicing events regulated by ABA or SR34a (see Fig. 6a), the 48 events changed by ABA exclusively in wild-type seeds were similarly affected by the *sr34a-1* mutation under control conditions (Fig. 6c, left). By contrast, the splicing changes for the 57 events affected by ABA only in the mutant showed no correlation with the changes imposed by the mutation alone (Fig. 6c, right), implying that SR34a does not repress these 57 events under control conditions. This indicates that the SR protein tends to prevent ABA-induced changes in splicing that are not already being repressed under basal conditions.

The 48 and 57 alternative splicing events ABA regulated specifically in either the wild type or *sr34a-1* (Fig. 6c) occurred respectively in 40 and 47 genes (Supplementary Data 11). A close look at the reported functions of the individual genes from these two groups revealed several ABA-related components (Supplementary Data 11), including seven genes for which reverse genetics has established a role in controlling sensitivity to ABA during early seedling development. Two of these, *RRP41L* (AT4G27490), which encodes a 3’-5’ exonuclease involved in cytosolic mRNA decay^45^, and *GRP7* (AT2G21660), encoding a glycine-rich RNA-binding protein^46^, were among the 40 genes differentially spliced in response to ABA only in wild-type seeds. In line with negative regulation of ABA responses, the other five genes reported to determine ABA sensitivity were among the 47 whose alternative splicing was changed by the hormone only in the *sr34a-1* mutant. They encoded three transcription factors — NF-YA6 (AT3G14020) which plays a role in seed development, seed germination and post-germinative growth^47^, AGL16 (AT3G57230) which controls the expression of stress-responsive genes^48^, and ABF4/AREB2 (AT3G19290), a master regulator of ABA signaling^49^ — as well as SIZ1 (AT5G60410), an E3 ligase that sumoylates key ABA signaling components such as the ABI5 transcription factor^50^, and GTG2 (AT4G27630), a GPCR-type G-protein that positively regulates the ABA pathway^51^.

The majority of the alternative splicing events regulated by ABA in the wild type (57%) and *sr34a-1* (65%) were in RNAs bound by SR34a (Fig. 6d, Supplementary Data 11), indicating that the SR protein regulates directly most of the splicing response to ABA. Except for *NF-YA6*, whose RNA was not bound by SR34a, and *GTG2*, for which no binding site was found in the exons flanking the relevant ABA-regulated event, this was the case for the five other genes known to control ABA sensitivity (Fig. 6e, f). Figure 6e illustrates those whose ABA-responsive splicing was mediated by SR34a, with an exonic iCLIP binding site identified in *RRP41L* immediately downstream of an ABA-mediated Alt3 event that produces a transcript with a longer 5’ UTR, while ABA regulation of an Alt5 event in the second exon of *GRP7* was associated with SR34a binding at the exons flanking the regulated alternative sequence. Among the genes whose ABA splicing regulation was repressed by SR34a (Fig. 6f), an IR event in *ABF4/AREB2* associated with strong SR34a binding at the flanking exonic sequences, inhibiting ABA retention of this intron. Furthermore, SR34a repression of an ABA-induced Alt5 event at the end of the *SIZ1* coding sequence was linked to increased binding of the splicing factor both within the upstream exon and the intronic region immediately downstream of the alternative sequence (Fig. 6f). Finally, ABA-mediated exclusion of an intron in the *AGL16* RNA (Fig. 6f) was associated with predominant exonic SR34a binding in the vicinity of the alternatively-spliced sequence.

In conclusion, our results indicate that the SR34a protein massively impacts the alternative splicing response to ABA, globally repressing the splicing changes induced by the hormone. In response to ABA, SR34a predominantly prevents alternative splicing changes that it does not already repress under basal conditions and directly targets most of these transcripts, several of which encode established key players of the ABA response.

## Discussion

Alternative splicing is a powerful and versatile means of gene regulation that has emerged as an important player in ABA mediation of plant responses to abiotic stress. Yet, little is known of the upstream molecular components that regulate this posttranscriptional mechanism to confer stress tolerance and of its role during the early stages of plant growth. Our study provides key insight into how a hitherto uncharacterized *Arabidopsis* SR protein regulates ABA responsive alternative splicing and shapes stress responses mediated by the phytohormone during seedling establishment. After reverse genetics revealed that the SR34a protein negatively regulates the ABA-mediated arrest of early plant development, promoting the transition to autotrophic growth under osmotic stress, we combined RNA-seq and iCLIP to identify the RNA molecules and sequences targeted by this SR protein to control the ABA transcriptomic and posttranscriptomic response during seedling establishment.

The first indication that the SR34a protein was involved in controlling stress responses mediated by the ABA phytohormone came from our finding that the *sr34a-1* loss-of-function mutant is hypersensitive to ABA and osmotic stresses during early seedling development. In agreement with this phenotype, RNA-seq analyses showed that the SR34a protein globally downregulates ABA-inducible gene expression under both control and ABA conditions, and represses alternative splicing changes induced by the hormone in numerous transcripts. Notably, our analysis revealed a lower number of DEGs than DAS events. Previous studies found that mutations in the *Arabidopsis SCL30a*^24^ or in both *SC35* and the entire *SCL* subfamily^13^ affected gene expression more than alternative splicing, while mutations in the rice *RS33*^18^ or the *Arabidopsis SR45a*^17^ genes predominantly affected alternative splicing,. These disparities are likely to reflect different individual SR protein contributions to distinct layers of gene regulation. The higher number of SR34a-regulated alternative splicing events suggests a preponderant role as a splicing regulator, though we cannot exclude that the protein fulfills broader functions in RNA metabolism, as described for many animal SR proteins^9^. In fact, fluorescent loss in photobleaching (FLIP) experiments performed by Stankovic et al.^29^ indicate that the SR34a protein shuttles between the nucleus and the cytoplasm, pointing to potential roles in post-splicing processes such as NMD or translation.

Though extensively reported in animal systems, the identification of the RNA sequences bound by plant SR proteins remains a challenge. Using RIP-seq, the transcripts associated with the *Arabidopsis* SR45 protein in inflorescences^17^ or 10-day-old seedlings^20^ were found to be enriched in very different sequence motifs. While this discrepancy is likely to reflect tissue specificity, it could also stem from RNAs associated to other proteins in the SR45 complex due to formaldehyde-induced protein-protein crosslinking^20^. Another limitation of the RIP-seq approach is that it does not identify binding sites directly, merely inferring their sequence through alignment of the retrieved transcripts. Here, we succeeded in adapting the iCLIP2 protocol to the challenging *Arabidopsis* seed tissue. We were thus able to map SR34a crosslinks with high precision and discovered that the SR protein binds preferentially exons in the vicinity of splice sites, consistent with the reported association of animal SR proteins with exonic splicing enhancers (ESEs)^52^. Importantly, our study revealed that SR34a crosslinks preferentially to two motifs, GCUs and U-rich sequences. While previous work has indicated that several other *Arabidopsis* splicing factors, including SR45^20^, RZ-1C^36^ and GRP7^31^, bind U-rich motifs, the observed enrichment in these sequences could result from an UV-C-mediated crosslinking bias towards uridines^42^. The more striking binding to GCU-rich sequences suggests that this motif is an actual ESE recognized by SR34a. A computational prediction analysis in *Arabidopsis* detected GCU triplets among the top-scored 9-mer ESEs and confirmed them experimentally^53^, but no splicing factor recognizing this sequence has been identified to date. The findings reported here support the relevance of this motif and that of SR34a in promoting splice site recognition. Our identification of the GCU sequence contrasts not only with the GA-rich motifs reported for the mammalian SRSF1^54,55^, pointing to divergent RNA-recognition mechanisms in plant and animal SR proteins, but also with the GAAG-containing motif found both to bind the *Arabidopsis* SCL30^14^ and SCL33^22^ in vitro and to be enriched in the intron-flanking regions of SR45-associated transcripts identified by RIP-seq^20^. This points to distinct sequence specificities among plant SR proteins, as already established among their metazoan counterparts^41,54,56^.

In addition to extensive transcriptional changes, the ABA phytohormone has been shown to significantly affect alternative splicing patterns in *Arabidopsis* seedlings^43,57–59^ as shown here for germinated seeds. Our observation that ABA alters alternative splicing in genes that are not changed by the hormone at the expression level supports the notion that transcription and splicing are independent regulatory layers in embryonic tissues, as already reported for later developmental stages^43,58–63^. Consistent with this, recent profiling of the temporal dynamics of ABA-responsive gene expression and alternative splicing revealed a greater contribution of the latter process at early time points^58^, supporting an autonomous role for splicing regulation in the modulation of ABA responses.

The molecular components that link ABA signaling and alternative splicing to achieve plant stress tolerance remain elusive. Given that the expression and splicing patterns of *SR34a* are unaffected by ABA in germinated seeds (data not shown), posttranslational regulation by the phytohormone appears likely. In fact, SR proteins are subjected to extensive phosphorylation in their RS domain^5,9^, ABA-triggered stress signaling causes differential phosphorylation of several plant splicing factors including SR34a^65^, and ABA was recently reported to dephosphorylate SR45 to negative regulate ABA responses during early *Arabidopsis* growth^27^. While our confocal microscopy analyses did not reveal evident ABA-induced changes in the subcellular localization of SR34a, whether alterations in the phosphorylation status of the SR protein play a role in regulating its RNA-binding or shuttling activity to modulate stress signaling remains to be investigated.

Global analysis of the alternative splicing RNA-seq data revealed a close interconnection between the action of the SR protein and of the phytohormone during postgerminative growth (Fig. 7). Indeed, we observed a clear correlation between the individual effects of the *sr34a-1* mutation and the phytohormone on alternative splicing, i.e, loss of SR34a function tends to induce ABA-responsive alternative splicing in the absence of the hormone. Given that the majority of the DAS events were associated with SR34a binding near the regulated sequences, this correlation implies that the SR protein attenuates ABA signaling already under basal conditions by directly repressing ABA-inducible alternative splicing changes. In agreement, the mutant exhibits both derepression of ABA-inducible gene expression and delayed cotyledon greening under control conditions.

**Fig. 7.**
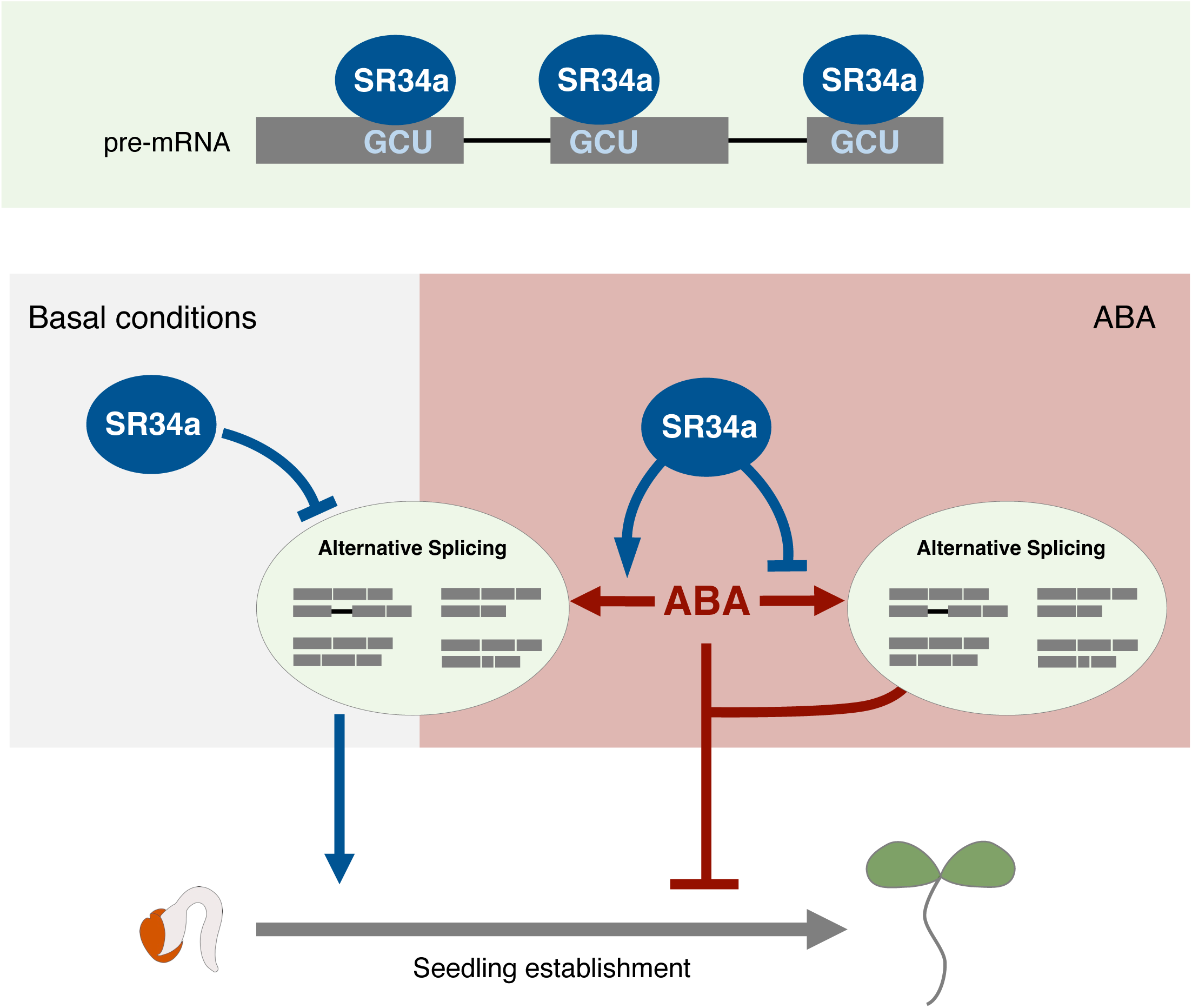
Model of SR34a control of ABA-responsive alternative splicing during early seedling development. In germinated seeds**, t**he SR34a splicing factor binds exonic sequences enriched in GCU motifs near 5’ and 3’ splicing sites to regulate all types of alternative splicing events under both basal and ABA conditions. Under basal conditions, SR34a represses ABA-inducible alternative splicing changes, thus promoting seedling establishment, as corroborated by delayed cotyledon greening observed in the *sr34a-1* mutant. Upon exposure to ABA, SR34a plays a dual role in the mediation of the alternative splicing response — while it mediates ABA-induced changes in a set of alternative splicing of events that it largely represses under basal conditions, SR34a represses ABA-induced changes in another larger set of alternative splicing events, thus counteracting ABA inhibition of postgermination growth, as corroborated by the ABA hypersensitivity of *sr34a-1*. Importantly, several of the ABA-responsive alternative splicing events under SR34a control occur in genes encoding established determinants of plant ABA sensitivity (e.g. *GRP7*, *RRP41L, ABF4*, *SIZ1*, *AGL16)*.

The stronger physiological phenotype displayed by the *sr34a-1* mutant under ABA and osmotic stress when compared to control conditions indicates a prevalent function for SR34a in the plant’s response to ABA-mediated stress. Our results demonstrate that this is achieved chiefly through posttranscriptional control, as loss of function of the RNA-binding protein markedly disturbed the ABA-induced alternative splicing response in germinated seeds while barely affecting ABA responsive gene expression, again corroborating a predominant role for SR34a in controlling splicing rather than gene expression levels. Of the alternative splicing events found to be changed by ABA exposure, the nearly half that were regulated specifically in wild-type seeds, i.e. those mediated by SR34a, showed the globally observed tendency to be downregulated by the SR protein under control conditions. Remarkably, the majority of ABA-responsive events that were affected specifically in the *sr34a-1* mutant, i.e. those derepressed by the mutation, were in general not regulated by SR34a in the absence of the hormone. Thus, the SR34a protein counteracts ABA-mediated changes in alternative splicing, not only in response to but also in anticipation of hormone exposure, underscoring a meaningful role in preventing the ABA splicing response in germinated seeds.

Most of the alternative events regulated by ABA in germinated seeds were associated with direct binding of SR34a. Notably, these SR34a-bound transcripts included five known determinants of ABA sensitivity during early seedling development, which are thus likely to mediate the physiological function of SR34a. While two of these transcripts, namely *GRP7*^46^ and *RRP41L*^45^, were found to require a functional SR34a protein for their ABA splicing regulation, we also found that the SR protein binds to and represses ABA-mediated alternative splicing of *ABF4*^49,65^, *AGL16*^48^ and *SIZ1*^50^. This is in line with the strong ABA hypersensitivity of the *sr34a-1* mutant and its global upregulation of ABA-inducible genes. Intriguingly, we found no substantial effect of the splicing factor on the molecular ABA responsiveness of the *ABI3* and *ABI5* genes, encoding recognized players in the ABA-induced postgermination growth arrest^2,4^. This suggests either that SR34a control of this developmental arrest is independent from these two components or that its splicing function indirectly affects their activity, for instance via SR34a repression of ABA-mediated splicing changes in *SIZ1*, which attenuates ABA signaling through posttranslational regulation and subsequent degradation of ABI5^50^.

While providing novel mechanistic insight into the function of plant SR proteins, the present study underlines the importance of alternative splicing regulation in modulating ABA control of postgerminative growth. Our data are fully consistent with SR34a counteracting ABA inhibition of seedling establishment by binding exonic elements near 5’ and 3’ splice sites to repress specific ABA-responsive alternative splicing events. In addition to expected changes in protein function or transcript stability/translation stemming for example from direct repression of splicing in the coding sequences of *SIZ1* and *AGL16* or the 5’UTR of *ABF4*, SR34a function presumably also relies on its numerous other target genes, which may fulfil yet-uncharacterized ABA functions. Detailed genetic analyses will be required to disclose the individual contribution of the multiple splicing targets of SR34a to its relief of ABA inhibition of seedling establishment.

## Methods

### Plant material and growth conditions

The *Arabidopsis thaliana* ecotype Columbia (Col-0) was used as the wild type in all experiments. PCR-based genotyping of the GK-803A10 line (obtained from NASC) with primers specific for *SR34a* and the left border of the T-DNA (Supplementary Data 12) followed by sequencing of the DNA/T-DNA junction confirmed the insertion site and allowed isolation of the *sr34a-1* homozygous line.

Seeds were surface-sterilized for 10 min in 50% (v/v) bleach and 0.07% (v/v) TWEEN^®^20, stratified for 3 days at 4°C in the dark and plated on MS medium [1X Murashige and Skoog (MS) salts (Duchefa Biochemie), 2.5 mM MES (pH 5.7), 0.5 mM myo-inositol and 0.8% (w/v) agar], before transfer to a growth chamber under 16-h photoperiod (long-day conditions) or continuous light (cool white fluorescent bulbs, 18W840, 4,000 K and 100 µmol m^-2^ s^-1^) at 22°C (light period) or 18°C (dark period) and 60% relative humidity. After 2-3 weeks, seedlings were transferred to soil in individual pots for seed production.

For all assays, fully mature siliques from dehydrated plants were collected and stored in the dark for at least 3 weeks at room temperature. For seed germination and cotyledon greening assays, 80-100 seeds per biological replicate were sown on MS medium supplemented or not with the appropriate concentrations of NaCl, mannitol or ABA (Duchefa Biochemie; A0941) and then transferred to long-day conditions, except for the determination of seed germination and cotyledon greening rates under control conditions, which was conducted under continuous light to avoid the effect of long dark periods during a short experimental time frame. Percentages of seed germination, defined as protrusion of the radicle through the seed coat, and cotyledon greening, defined as the presence of green cotyledons, were calculated over the total number of plated seeds. The results presented are representative of at least three independent experiments.

For RT-qPCR analyses, seeds were surface-sterilized, stratified, sown on Whatman^®^ paper placed on top of MS agar medium and transferred to continuous light conditions. For the RT-qPCR in Figure 1e as well as for the RNA-sequencing, SR34a-GFP immunoprecipitation and iCLIP2 assays, seeds were transferred after 44 h to MS (control conditions) or ABA-containing MS media for 2 h before UV-crosslinking (iCLIP2) or direct sample collection.

### Generation of transgenic lines

To generate the *SR34a:SR34a-GFP* construct, a 4142-bp PCR fragment containing the genomic *SR34a* sequence and including the endogenous regulatory elements from 1980 bp upstream of the translation start codon was subcloned into the pGEM-T Easy vector and introduced into an eGFP-tagged version of the binary pBA002 vector using the SacI/AatII restriction sites. In addition, a 374-bp PCR fragment corresponding to the predicted 3’ untranslated region (UTR) of *SR34a* was cloned downstream of the *eGFP* using the SpeI/SacI restriction sites. Transformation of this construct into *sr34a-1* mutant plants was achieved by the floral dip method^66^ using *Agrobacterium tumefaciens* strain GV3101. Homozygous *SR34a:SR34a-GFP_sr34a-1* complementation lines were obtained from self-pollination of the previously generated hemizygous lines. Homozygous plants overexpressing the *eGFP* alone were also generated and isolated after transformation of the *sr34a-1* mutant with the *pro35S:eGFP* construct.

### Expression and alternative splicing analysis of individual genes

Total RNA was extracted from frozen ground tissues using the innuPREP Plant RNA kit (Analytic Jena) and digested with RQ1 DNase (Promega). First-strand cDNA synthesis was conducted on 1 µg RNA using the SuperScript III Reverse Transcriptase (Invitrogen) and an oligo-dT primer.

qPCR was performed using an ABI QuantStudio sequence detection system (Applied Biosystems) and Luminaris Color HiGreen qPCR Master Mix (Thermo Scientific) on 2.5 µL of cDNA (diluted 1:10) per 10-µL reaction volume containing 300 nM of each gene-specific primer (Supplementary Data 12). Reaction cycles were 95°C for 2 min (1X), 95°C for 30 s/ 60°C for 30 s/72°C for 30 s (40X), followed by a melting curve step to confirm the specificity of the amplified products. *UBQ10* and *ROC5* were used as reference genes. Each experiment was replicated at least three times.

For the alternative splicing analyses shown in Supplementary Figure 4, PCR with the NZYTaq II 2x Green Master Mix (NZYtech) was performed on cDNA from 3-4 biological replicates of germinated seeds (44 h after stratification, continuous light) using primers flanking the alternatively-spliced sequence (Supplementary Data 12) obtained from PASTDB (pastdb.crg.edu). Reaction cycles were 95°C for 3 min (1X), 95°C for 30 s/58°C for 30 s/72°C for 5 min (35X). The PCR products were then loaded on a 2% agarose gel and gel bands quantified using the ImageJ software (https://rsbweb.nih.gov/ij). The percentage of inclusion (PSI; Percent of Spliced In) for each alternative splicing event was calculated after quantification of the variants including (I) or excluding (S) the given alternative sequence, with PSI = I/(I+S).

### Subcellular localization analysis

Confocal images of the SR34a-GFP fusion protein were obtained from 44-h germinated seeds of the *SR34a:SR34a-GFP_sr34a-1* transgenic line, transferred to MS (control conditions) or MS supplemented with 0.5 µM ABA (Duchefa Biochemie; A0941) media for 2 h, using a Zeiss LSM980 laser scanning microscope. Excitation wavelength used to visualize the fluorescence from GFP was 488 nm, while the detection wavelength was set at 550 nm.

### Protein immunoprecipitation and immunoblot analyses

For immunoprecipitation of SR34a-GFP or GFP alone, an initial amount of approximately 50 mg of *SR34a:SR34a-GFP_sr34a-1* seeds was used. After grinding of the plant material in liquid N_2_, 800 µL of cell lysis buffer [50 mM Tris-HCl (pH 7.5), 150 mM NaCl, 4 mM MgCl_2_, 0.25% Igepal CA-630, 1% SDS, 0.25% sodium deoxycholate, 5 mM DTT, Complete Protease Inhibitor (Roche), 100 U/mL RiboLock (Thermo Fisher), 1 mM phenylmethylsulfonyl fluoride]^31^ were added to the powder. Lysates were precleared with 50 µL of sepharose beads for 1 h at 4°C with constant rotation and subjected to immunoprecipitation with 25 µL GFP Trap agarose beads (Chromotek, gta-20) during 1 h at 4°C with constant rotation. The beads were then washed four times with 1 mL of cooled RIP-washing buffer [50 mM Tri<u>s</u>-HCl (pH 7.5), 500 mM NaCl, 4 mM MgCl_2_, 0.5% Igepal CA-630, 1% SDS, 0.5% sodium deoxycholate, 2 M urea, 2 mM DTT, Complete Protease Inhibitor (Roche)]. The proteins were eluted from the beads in 20 µL 2X Laemmli buffer for 5 min at 95°C, resolved on 8% SDS polyacrylamide gels and transferred to PVDF membranes (Immoblon-P, Millipore), which were then blocked with 5% non-fat dry milk for 2 h. The membranes were probed overnight at 4°C with monoclonal anti-GFP primary antibodies (Roche, 11814460001) at 1:1,000 and then with anti-mouse peroxidase-conjugated secondary antibodies (Jackson ImmunoResearch, 115-035-146) at 1:20,000 for 2 h at room temperature. The peroxidase activity associated with the membrane was visualized by chemiluminescence.

### RNA-seq sample preparation and sequencing

Approximately 50 mg of Col-0 wild-type and *sr34a-1* mutant seeds (3 biological replicates) were stratified and sown on Whatman^®^ paper placed on top of MS medium. Seeds were germinated for 44 h and then transferred to MS (control) or MS supplemented with 0.5 µM ABA (Duchefa Biochemie; A0941) media for 2 h. Total RNA was extracted using the innuPREP Plant RNA Kit (Analytic Jena). The RNA-seq libraries were prepared and sequenced by Novogene (UK) Co. Ltd using the Novaseq 6000 system (Illumina, Inc) with a read length of 2 x 150 bp.

### iCLIP2-seq sample preparation and sequencing

UV-crosslinking and immunoprecipitation protocols established for mammalian cells, *Caenorhabditis elegans* and *Arabidopsis* seedlings^31,36–38^ were adapted to our plant experimental conditions. After stratification, 3 g of *SR34:SR34a-GFP_sr34a-1* or *35S:GFP_sr34a-1* seeds were germinated for 44 h before transfer to MS (control conditions) or MS supplemented with 0.5 µM ABA (Duchefa Biochemie; A0941) media for 2 h. Germinated seeds were then irradiated with 254-nm UV light at 2,000 mJ/cm^2^ using an UVC-515 Ultraviolet Multilinker, frozen in liquid N_2_ and ground to a homogeneous powder. For each sample, the resulting powder was equally distributed in 3 50-mL Falcon tubes, and 25 mL of pre-heated cell lysis buffer [50 mM Tris-HCl (pH 7.5), 150 mM NaCl, 4 mM MgCl_2_, 0.25% Igepal CA-630, 1% SDS, 0.25% sodium deoxycholate, 5 mM DTT, Complete Protease Inhibitor (Roche), 100 U/mL RiboLock (Thermo Fisher), 1 mM phenylmethylsulfonyl fluoride] were added to each tube. Lysates were precleared with 200 µL of sepharose beads for 1 h at 4°C under constant rotation and then subjected to immunoprecipitation with 50 µL of GFP Trap agarose beads (Chromotek, gta-20) for 2 h at 4°C under constant rotation. The beads were washed 4 times with 1 mL of cooled RIP-washing buffer [50 mM Tris-HCl (pH 7.5), 500 mM NaCl, 4 mM MgCl_2_, 0.5% Igepal CA-630, 1% SDS, 0.5% sodium deoxycholate, 2 M urea, 2 mM DTT, Complete Protease Inhibitor (Roche)] and twice with 1 mL of cooled original iCLIP wash buffer [20 mM Tris-HCl (pH 7.4], 10 mM MgCl_2_, 0.2% TWEEN^®^20]^30^. The immunoprecipitate was next treated with 2 µL of Turbo DNase (Thermo Fisher) for 10 min at 37°C. For on-bead RNase digestion, 10 U of RNase I (Thermo Fisher) were added together with the DNase. For library preparation, the RNAs were dephosphorylated and the L3 linker (Supplementary Data 1) ligated to the 3’ ends using RNA ligase (NEB). The RNA 5’ ends were radiolabeled using [γ-^32^P]-ATP and polynucleotide kinase and the covalently-linked protein-RNA complexes separated on a 4-12% NuPAGE Bis-Tris gel (Thermo Scientific) before transfer onto a nitrocellulose membrane. The RNA-protein complexes were revealed by autoradiography before the regions above the fusion protein were cut out and subjected to proteinase K treatment, leaving a polypeptide at the interaction site. The RNA was then isolated from the membrane using phenol-chloroform and concentrated with the RNA Clean and Concentrator kit (Zymo Research, R1013), followed by reverse transcription using the RT Oligo primer (Supplementary Data 12 and SuperScript III. cDNAs were subsequently purified using the DynaBeads MyONE Silane system (Thermo Fisher, 37002D), and a second adapter L#clip2.0 (Supplementary Data 12) containing a 6-nt barcode and a 9-nt unique molecular identifier sequence^30^ was ligated to the 3’ end of the cDNA. Each sample was ligated with a different L#clip2.0 adapter (see Supplementary Data 12) and purified again with the DynaBeads MyONE Silane kit (Thermo Fisher, 37002D). cDNAs were PCR pre-amplified — 98°C for 30 s (1X), 98°C for 10 s/65°C for 30 s/72°C for 30 s (6X), 72°C for 3 min (1X) — using the 2X Phusion HF PCR MasterMix (Thermo Fisher) and P5Solexa_s and P3Solexa_s primers (Supplementary Data 12), with primer dimers being removed using the ProNex Size-Selective Purification System (Promega, NG2001). After a second PCR amplification— 98°C for 30 s (1X), 98°C for 10 s/65°C for 30 s/72°C for 30 s (10X), 72°C for 3 min (1X) — using the 2X Phusion HF PCR MasterMix (Thermo Fisher) and P5Solexa and P3Solexa primers (Supplementary Data 12) on the whole eluted volume and a second ProNEX purification step (Promega, NG2001) to remove residual primers, RNA concentrations and the median size of the RNA fragments from the libraries were assessed using a 5200 Fragment Analyzer System (Agilent). Finally, the libraries were diluted to 10 nM and submitted to high-throughput sequencing using the NextSeq 500 system (Illumina) at the IGC Genomics Unit. Three biological replicates from *SR34a:SR34a-GFP_sr34a-1* germinated seeds subjected to either control or ABA conditions as well as 3 biological replicates from a pool of control and ABA-treated *35S:GFP_sr34a-1* germinated seeds (negative control) were used, resulting in the sequencing of a total of 9 libraries.

### RNA-seq identification and quantification of differential gene expression

Quantification of total mRNA levels was performed using *vast-tools* v2.5.1^67^. This tool provides the corrected-for-mappability RPKMs (cRPKMs), corresponding to the number of mapped reads per million mapped reads divided by the number of uniquely mappable positions of the transcript^68^. To identify differentially-expressed genes (DEGs) between wild-type and *sr34a-1* germinated seeds or between control and ABA conditions, *vast-tools_compare_expr* command was employed with the option -norm, which allows a quantile normalization of cRPKMs between samples. Moreover, genes with read counts < 50 and that were not expressed at cRPKM > 5 across all replicates of at least one of the two samples compared were filtered out. Finally, those genes with a fold change of at least 2 between each of the individual replicates from each sample analyzed were defined as DEGs (Supplementary Data 1, 3).

### RNA-seq identification and quantification of differential alternative splicing

To quantify alternative splicing from each individual sample we used *Vast-tools* v2.5.1^67^, which maps the RNA-seq data to the araTha10 library, based on Ensembl Plants v31 and composed of an extended annotation with all exon-exon and exon-intron junction sequences found in the *A. thaliana* genome^70^. The *vast-tools* program then quantifies intron retention (IR), exon skipping (ES), alternative 5’ splice site (Alt5) and alternative 3’ splice site (Alt3) events in each sample and provides the percentage of inclusion (PSI) of the putative alternative sequence using only exon-exon (or exon-intron for IR) junction reads. The tool also provides information on the read coverage that sustains quantification of the PSI (see https://hub.com/vastgroup/vast-tools for details). The *vast-tools compare* command was then used to define differential alternative splicing (DAS) events in the *sr34a-1* mutant or in response to ABA. To evaluate more and less sequence inclusion of all Alt5 and Alt3 events, not only of the most external splice sites, we used the –legacy_ALT *vast-tools compare* option. Moreover, the –min_ALT_use 25 option was used to ensure that Alt3 and Alt5 events were located in exons with a minimum PSI of 25 in each compared sample. Finally, splicing events with a |ΔPSI| > 15 between the compared pair of samples and whose PSI distribution did not overlap (–min_range 5) were defined as DAS events (Supplementary Data 5, 11). Furthermore, non-regulated alternative splicing events (Supplementary Data 9, 10) were defined as those with a 10 ≤ PSI ≤ 90 in at least one sample (e.g. wild type or *sr34a-1*), and a |ΔPSI| < 5 between the two compared samples.

### Gene ontology enrichment analyses and predicted protein impact of differential alternative splicing

To identify significantly enriched biological processes, molecular functions and cellular components among the different sets of genes, analyses were performed using the functional annotation classification system DAVID^71^. For each comparison, only genes with transcripts or events that passed equivalent filters than those used to define DEGs or DAS events were used as a background.

The gene location and ORF predictions for all DAS events were retrieved from the Downloads section of PastDB (http://pastdb.crg.eu/wiki/Downloads). Briefly, this tool classifies the alternative sequences based on their location within the gene structure and determines whether the sequence preserves the reading frame or contains in-frame stop codons predicted to trigger RNA-mediated nonsense mediated decay (NMD) (see ref. 43 for details). These criteria generate three different groups: DAS events occurring in the untranslated regions (UTRs), potentially affecting protein levels (by including or excluding a premature termination codon) or potentially producing alternative protein isoforms.

### iCLIP2-seq read processing and peak calling

Illumina iCLIP reads were quality checked with FastQC (0.11.9) and the sample barcode distribution with awk (GNU awk 5.0.1). Reads were demultiplexed and sequencing adapters removed using Flexbar (3.5.0). After a second quality check with FastQC, reads were quality- and length-trimmed with Flexbar. A genome index was created using STAR (2.7.3a) with the *A. thaliana* genome version TAIR10 and gene annotation from Araport (version 11). Reads were then mapped using STAR and the created genome index, allowing only softclipping of 3’ ends (--alignEndsType Extend5pOfRead1). PCR duplicates were removed using umi_tools (1.1.4) by considering the UMI tag in the read-ID and the mapping coordinates. Around 7-12 million and 46,000-120,000 uniquely mapped reads per replicate were identified in the SR34a-GFP and GFP samples, respectively (Supplementary Data 13). The uniquely mapped and deduplicated reads from each replicate were merged and peak called with PureCLIP (1.3.1) in standard mode. Peak clusters (directly adjacent peaks) were reduced to the peak with the highest reported PureCLIP score. The peak coordinates (1-nt width) were transformed into binding sites by extending them with 4 nt (−4…0…+4) in both directions using bedtools (2.30.0). Binding sites with only a single crosslinked position (1 out of 9) were considered artifacts and therefore removed. The reproducibility of binding sites was analyzed by overlapping the binding site locations with crosslinks from each replicate and the distribution used to determine a reproducibility threshold. The 30% quantile was chosen for both the mock- and ABA-treated SR34a samples. Binding sites from SR34a-GFP in control conditions were considered reproducible if they passed the 30% quantile threshold (at least 5, 4 or 4 crosslinks for replicate 1, 2 or 3) in at least 2 of 3 replicates. For the ABA-treated SR34a-GFP sample, the crosslink thresholds were set to 3, 5 and 4 crosslinks in 2 out of 3 replicates, respectively. The GFP control was not tested for reproducibility due to the comparably low amount of uniquely mapped reads. Reproducible binding sites of SR34a overlapping with binding sites from the GFP control were removed using bedtools. Tables listing the target transcripts (Supplementary Data 14, 15) were generated by overlapping coordinates of reproducible SR34a binding site locations with representative gene models from Araport (version 11). Crosslink tracks for visual inspection were generated from uniquely mapped read coordinates using bedtools. For motif discovery with STREME (5.4.1), the sequences of reproducible binding sites were extracted using bedtools in strand specific mode (-s).

## Data availability

The raw and processed RNA-seq and iCLIP data generated in this study have been deposited in the NCBI GEO database (https://www.ncbi.nlm.nih.gov/gds) under accession number GSE248563.

## Acknowledgments

We thank V. Nunes for excellent plant care, T. Paixão for valuable help with statistical analyses, and the Instituto Gulbenkian de Ciência Plant and Advanced Imaging Units for technical support. This work was funded by Fundação para a Ciência e a Tecnologia (FCT) through PhD Fellowship UI/BD/152253/2021 (to R.J.R.Y.), grant PTDC/ASP-PLA/2550/2021 and funding from the research unit GREEN-it “Bioresources for Sustainability” (https://doi.org/10.54499/UIDB/04551/2020), as well as by the German Research Foundation (DFG) through grant STA653/14-1 (to D.S). T.L. was supported by Marie Skłodowska-Curie Individual Fellowship MSCA-IF-2015 (project 706274) and G.M. by the Spanish Ministry of Science and Innovation through grants RYC2020-030160-I and PID2021-125223NA-I00.

## Author contributions

T.L., T.K., D.S. and P.D. designed the research, T.L. and R.J.R.Y. performed the experiments, and G.M. and M.L. conducted the computational analyses of the RNA-seq and iCLIP2 data, respectively. T.L. and P.D. wrote the manuscript and prepared the figures and tables. All authors contributed to the interpretation of results, critically reviewed the manuscript, and approved its final version.

## Competing interests

The authors declare no competing interests.

## Supplementary information

**Supplementary Fig. 1 ∣ Molecular characterization of the *sr34a-1* mutant. a** Schematic representation of the *SR34a* gene (boxes indicate exons with UTRs in gray, lines between boxes represent introns, and arrows indicate the location of *SR34a*-specific primers) and structure of the corresponding SR34a protein (RRM, RNA recognition motif; RS, arginine/serine-rich domain). The red asterisk marks the position of the predicted truncation in the *sr34a-1* mutant. **b** RT-qPCR analysis of *SR34a* transcript levels in Col-0 wild-type (WT, white bars) and *sr34a-1* mutant (black bars) seeds germinated for 44 h (means ± SE, *n* = 4), using primers upstream of (F1/R1) or flanking (F2/R2) the T-DNA insertion. Expression levels in the Col-0 WT were set to 1. Asterisks indicate statistically significant differences from the WT (*** p < 0.001; Student’s *t*-test).

**Supplementary Fig. 2 ∣ Characterization of transgenic *sr34a-1* complementation lines. a** RT-qPCR analysis of *SR34a* transcript levels in seeds from the Col-0 wild type (WT), the *sr34a-1* mutant and the *SR34a:SR34a-GFP_sr34a-1* C1 and C2 complementation lines germinated for 44 h (means ± SE, *n* = 3), using the F1/R1 primers shown in Supplementary Figure 1a. Expression levels in the Col-0 WT were set to 1. Asterisks indicate statistically significant differences from the WT (** p < 0.01; Student’s *t*-test). **b** Quantification of cotyledon greening percentages of seedlings from the Col-0 WT, the *sr34a-1* mutant and the C1 and C2 complementation lines scored 10 d after stratification and growth under different ABA concentrations. Asterisks indicate statistically significant differences from the WT (* p < 0.05; Student’s *t*-test). **c** Confocal laser scanning microscopy images of the SR34a-GFP protein from C1 transgenic seeds germinated for 44 h under control conditions (left image) and from the radicle of a C1 44-h germinated seed treated or not with ABA (right images). Scale bars: 100 µm (left image) or 50 µm (right images).

**Supplementary Fig. 3 ∣ ABA responsiveness of SR34a-regulated genes.** Expression levels of the genes up-(dark blue) or down-(light blue) regulated in the *sr34a-1* mutant under control (**a**) or ABA (**b**) conditions in wild-type (WT) samples under control or ABA conditions. Asterisks indicate statistically significant differences (*** p < 0.001; Mann & Whitney test). n.s., not significant.

**Supplementary Fig. 4 ∣ Validation of selected differential alternative splicing events detected by RNA-seq.** RT-PCR analysis of individual alternative splicing events found by RNA-seq to be differentially regulated between germinated Col-0 wild-type (WT) and *sr34a-1* mutant seeds in (**a**) AT2G25730 (event: AthINT0017262), (**b**) AT1G12750 (event: AthALTA0039695), (**c**) AT1G09140 (event: AthALTA0042857), (**d**) AT5G05550 (event: AthEX0010842) and (**e**) AT4G28680 (event: AthINT0109950). Arrowheads indicate the bands corresponding to each splice variant, and the alternatively-spliced sequence is shown in teal blue. RT-PCR graphs present Percent of Spliced In (PSI) values (means ± SE, *n* = 3-4) after quantification of the band intensities using the Image J software. Asterisks indicate statistically significant differences from the Col-0 WT (* p < 0.05, ** p < 0.01, *** p < 0.001; Student’s *t*-test).

**Supplementary Fig. 5 ∣ Identification of SR34a-GFP-RNA complexes in *Arabidopsis* germinated seeds. a** Immunoprecipitation of the SR34a-GFP fusion protein from *SR34a:SR34a-GFP_sr34a-1* C1 44-h germinated seeds treated or not with ABA and of the GFP protein from a pool of mock- and ABA-treated *35S:GFP_sr34a-1* 44-h germinated seeds. Lysates were subjected to immunoprecipitation with GFP Trap beads (IP+). Aliquots of the lysate (input, IN), the supernatant (SN) and IP+ were analyzed by immunoblotting with a-GFP antibodies. **b** Autoradiogram of RNA-protein complexes immunoprecipitated from *SR34a:SR34a-GFP_sr34a-1* C1 44-h germinated seeds after UV crosslinking (+UV) or no UV crosslinking (-UV). The signal comes from ^32^P-end-labeling of RNA. Treatment of the precipitate with RNase I (+ RNase) reveals the size of the SR34a-GFP protein + ∼5 kDa, corresponding to the short RNAs still bound by the protein after RNase treatment. The a-GFP immunoblot (bottom panel) identifies the precipitated SR34a-GFP protein.

**Supplementary Fig. 6 ∣ Autoradiograms of RNA-protein complexes isolated for iCLIP.** RNA-protein complexes immunoprecipitated from 44-h germinated seeds of the *SR34a:SR34a-GFP_sr34a-1* C1 (**a**) or *35S:GFP_sr34a-1* (**b**) transgenic lines treated or not with ABA. The rectangles indicate the regions that were cut out from the membrane for further RNA purification and sequencing.

**Supplementary Fig. 7 ∣ SR34a binding distribution and motifs under ABA conditions. a** SR34a iCLIP binding site peaks (i.e. middle position of binding sites) mapped in the vicinity of 5’ and 3’ splice sites under ABA conditions. **b** Binding motifs significantly enriched within SR34a binding sites identified under ABA conditions. **c** Number of iCLIP binding site peaks identified under ABA conditions as a function of their distance to GCU motifs in the *A. thaliana* genome (top), and number of GCU motif hits as a function of their distance to binding site peaks (bottom).

**Supplementary Fig. 8 ∣ Distribution and regulation of the different types of alternative splicing event under direct SR34a control.** Number of intron retention (IR), exon skipping (ES), alternative 5’ splice site (Alt5) and alternative 3’ splice site (Alt3) events among the 102 or 92 dDAS events in the *sr34a-1* mutant identified under control (gray) or ABA (red) conditions, respectively. The lighter and darker colored bars indicate the number of dDAS events in which the alternative sequence was respectively less or more included in the *sr34a-1* mutant.

## Notes

### Competing Interest Statement

The authors have declared no competing interest.

