## Supplementary Figures for "An *Arabidopsis* SR protein relieving ABA inhibition of seedling establishment represses ABA-responsive alternative splicing"

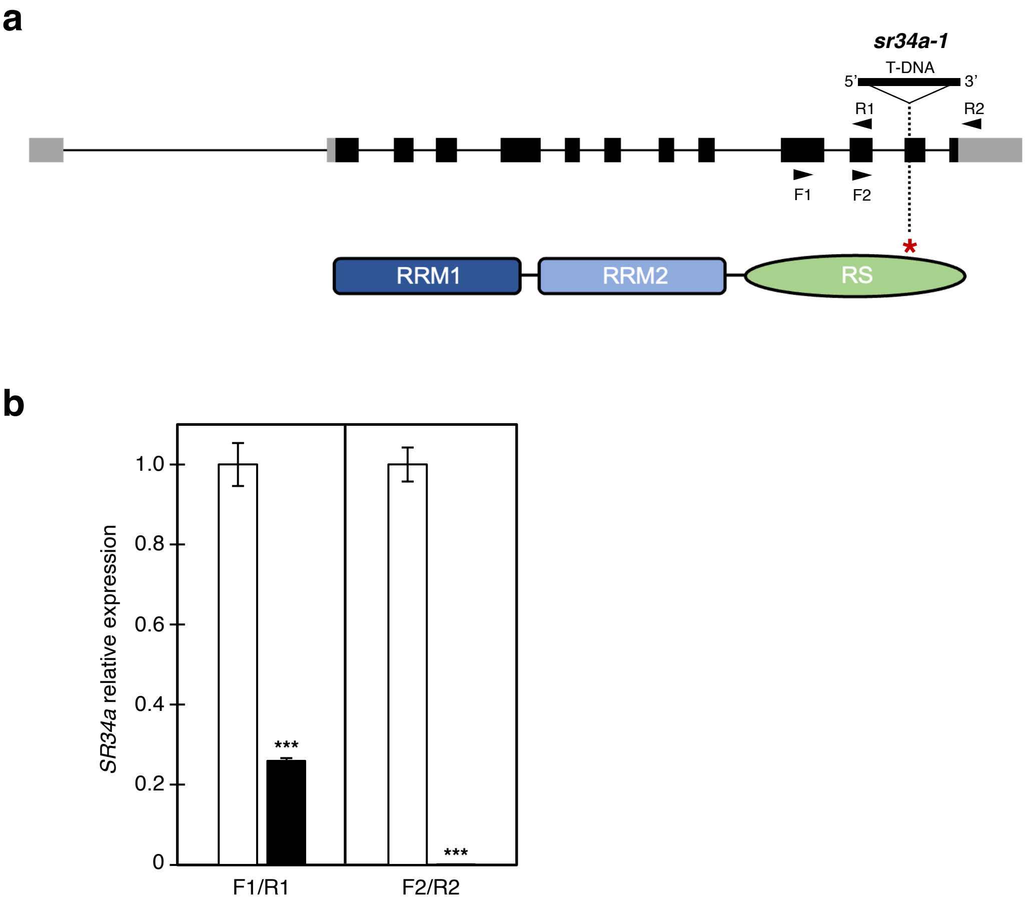


**Supplementary Fig. 1 | Molecular characterization of the *sr34a-1* mutant. a** Schematic representation of the *SR34a* gene (boxes indicate exons with UTRs in gray, lines between boxes represent introns, and arrows indicate the location of *SR34a*-specific primers) and structure of the corresponding SR34a protein (RRM, RNA recognition motif; RS, arginine/serine-rich domain). The red asterisk marks the position of the predicted truncation in the *sr34a-1* mutant. **b** RT-qPCR analysis of *SR34a* transcript levels in Col-0 wild-type (WT, white bars) and *sr34a-1* mutant (black bars) seeds germinated for 44 h (means ± SE, $n = 4$), using primers upstream of (F1/R1) or flanking (F2/R2) the T-DNA insertion. Expression levels in the Col-0 WT were set to 1. Asterisks indicate statistically significant differences from the WT (*** $p < 0.001$; Student's *t*-test).

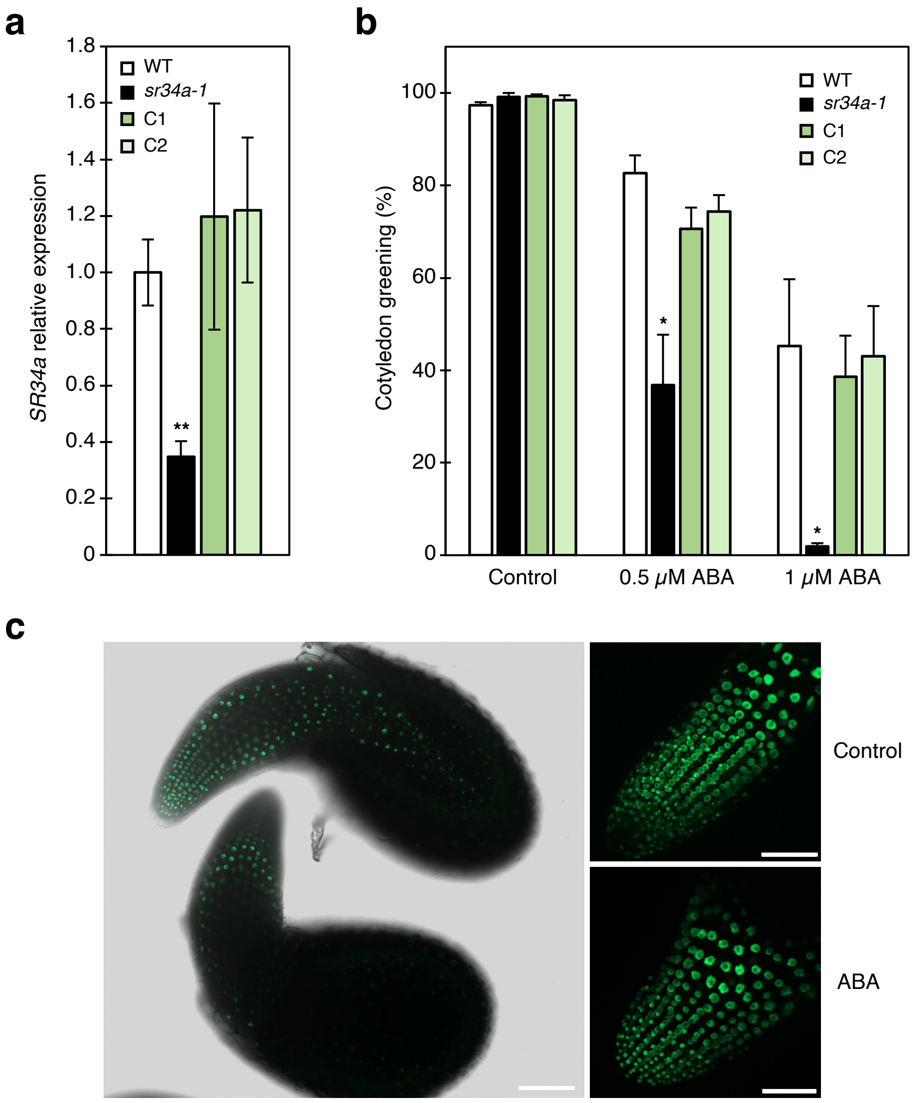


**Supplementary Fig. 2 | Characterization of transgenic *sr34a-1* complementation lines. a** RT-qPCR analysis of *SR34a* transcript levels in seeds from the Col-0 wild type (WT), the *sr34a-1* mutant and the *SR34a:SR34a-GFP_sr34a-1* C1 and C2 complementation lines germinated for 44 h (means ± SE, $n$ = 3), using the F1/R1 primers shown in Supplementary Figure 1a. Expression levels in the Col-0 WT were set to 1. Asterisks indicate statistically significant differences from the WT (** $p < 0.01$; Student's *t*-test). **b** Quantification of cotyledon greening percentages of seedlings from the Col-0 WT, the *sr34a-1* mutant and the C1 and C2 complementation lines scored 10 d after stratification and growth under different ABA concentrations. Asterisks indicate statistically significant differences from the WT (* $p < 0.05$; Student's *t*-test). **c** Confocal laser scanning microscopy images of the SR34a-GFP protein from C1 transgenic seeds germinated for 44 h under control conditions (left image) and from the radicle of a C1 44-h germinated seed treated or not with ABA (right images). Scale bars: 100 $\mu$m (left image) or 50 $\mu$m (right images).

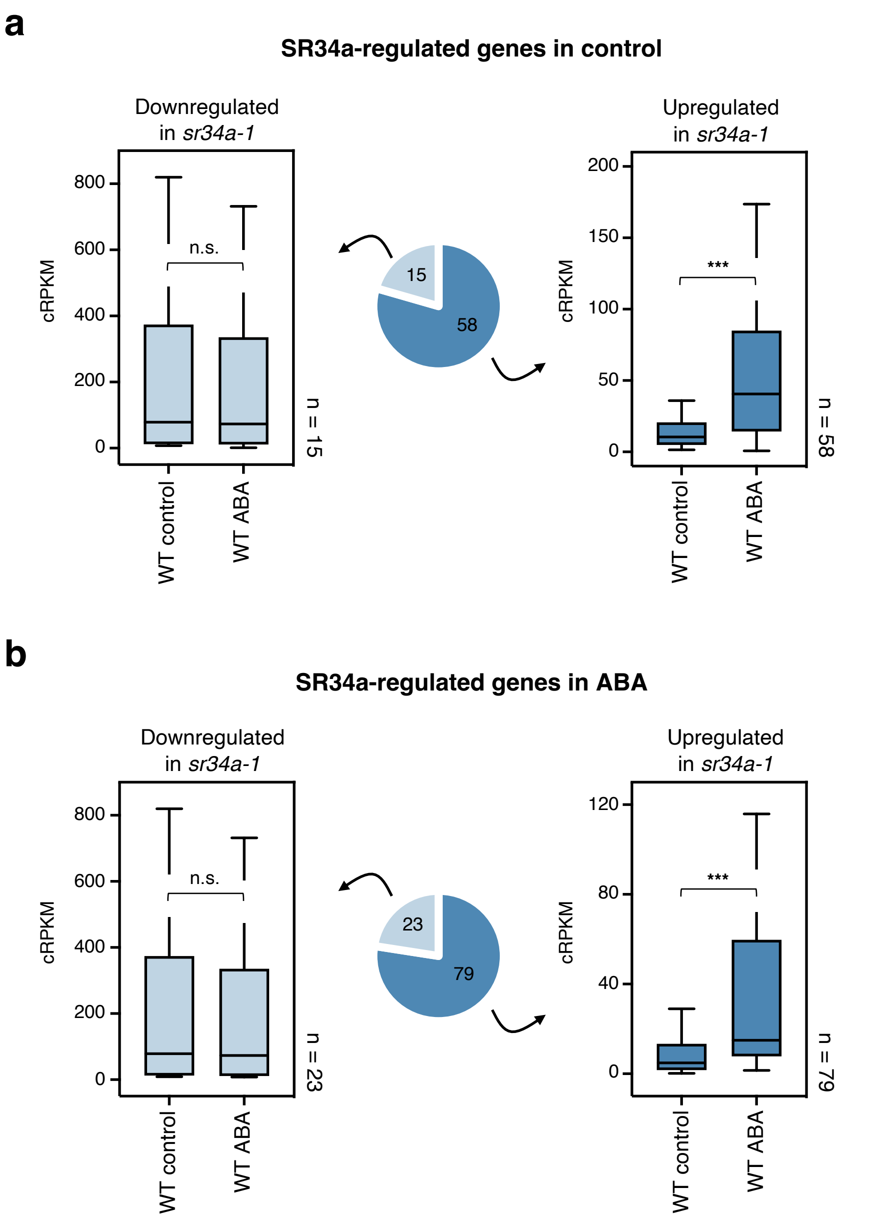


**Supplementary Fig. 3 | ABA responsiveness of SR34a-regulated genes.** Expression levels of the genes up- (dark blue) or down- (light blue) regulated in the *sr34a-1* mutant under control (**a**) or ABA (**b**) conditions in wild-type (WT) samples under control or ABA conditions. Asterisks indicate statistically significant differences (*** $p < 0.001$; Mann & Whitney test). n.s., not significant.

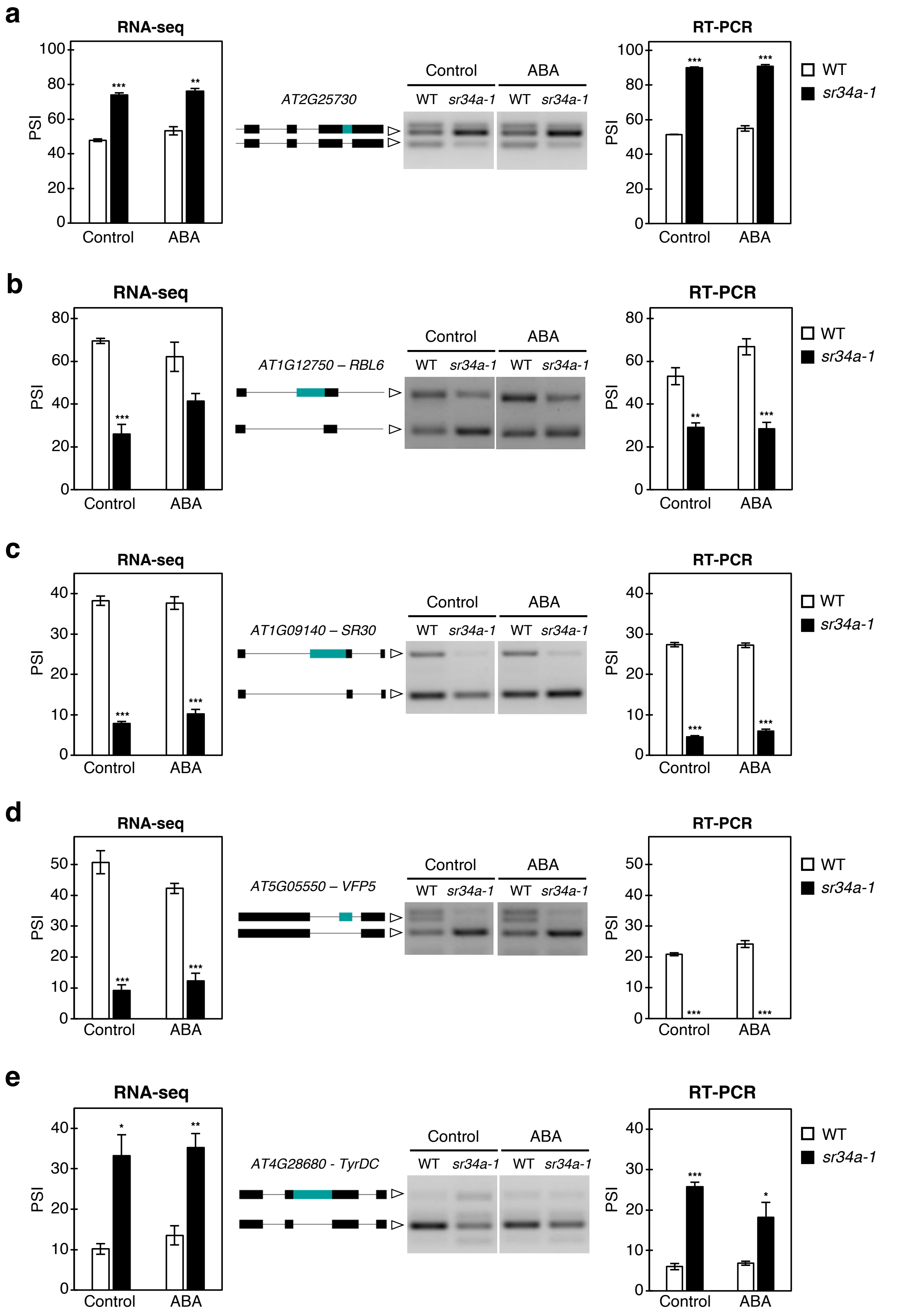


**Supplementary Fig. 4 | Validation of selected differential alternative splicing events detected by RNA-seq.** RT-PCR analysis of individual alternative splicing events found by RNA-seq to be differentially regulated between germinated Col-0 wild-type (WT) and *sr34a-1* mutant seeds in (**a**) AT2G25730 (event: AthINT0017262), (**b**) AT1G12750 (event: AthALTA0039695), (**c**) AT1G09140 (event: AthALTA0042857), (**d**) AT5G05550 (event: AthEX0010842) and (**e**) AT4G28680 (event: AthINT0109950). Arrowheads indicate the bands corresponding to each splice variant, and the alternatively-spliced sequence is shown in teal blue. RT-PCR graphs present Percent of Spliced In (PSI) values (means ± SE, $n$ = 3-4) after quantification of the band intensities using the Image J software. Asterisks indicate statistically significant differences from the Col-0 WT (* $p < 0.05$, ** $p < 0.01$, *** $p < 0.001$; Student's $t$-test).

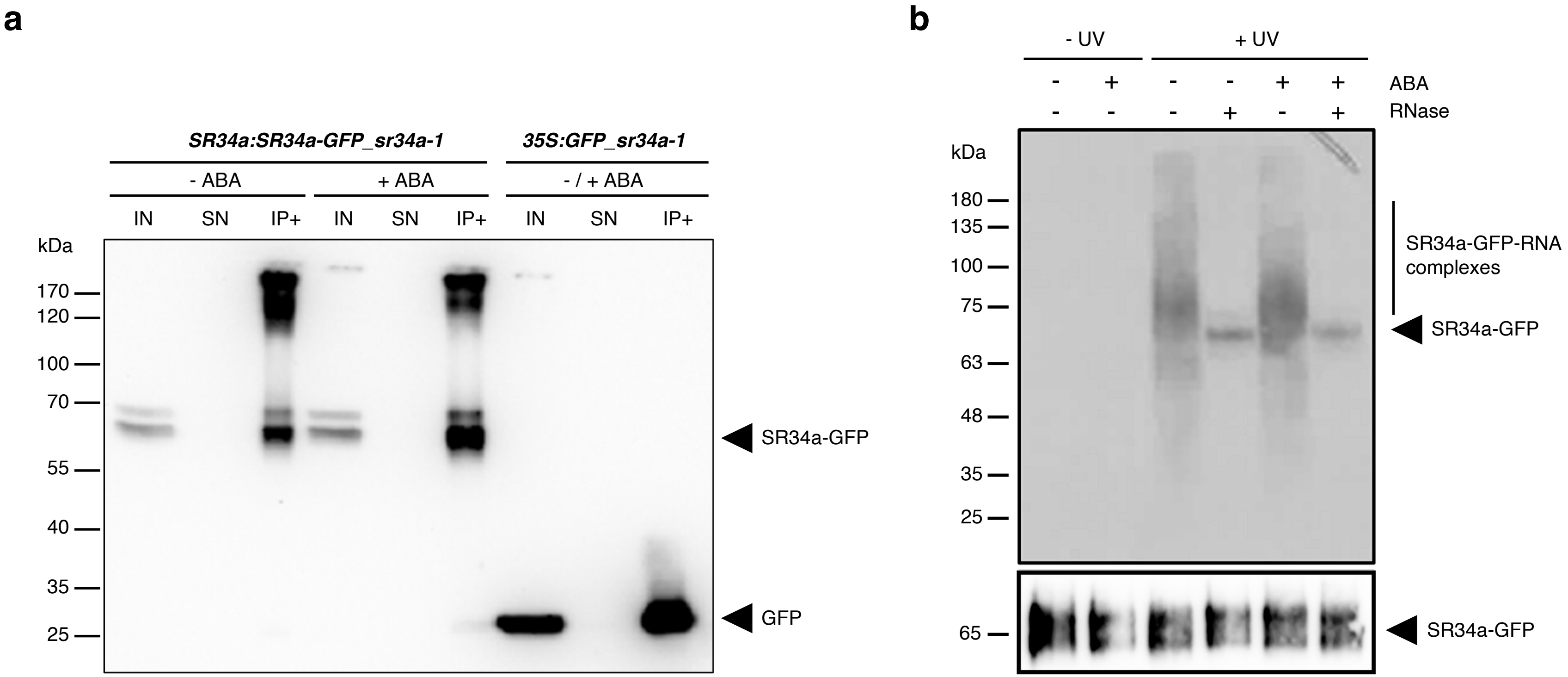


**Supplementary Fig. 5 | Identification of SR34a-GFP-RNA complexes in *Arabidopsis* germinated seeds. a** Immunoprecipitation of the SR34a-GFP fusion protein from *SR34a:SR34a-GFP_sr34a-1* C1 44-h germinated seeds treated or not with ABA and of the GFP protein from a pool of mock- and ABA-treated *35S:GFP_sr34a-1* 44-h germinated seeds. Lysates were subjected to immunoprecipitation with GFP Trap beads (IP+). Aliquots of the lysate (input, IN), the supernatant (SN) and IP+ were analyzed by immunoblotting with a-GFP antibodies. **b** Autoradiogram of RNA-protein complexes immunoprecipitated from *SR34a:SR34a-GFP_sr34a-1* C1 44-h germinated seeds after UV crosslinking (+UV) or no UV crosslinking (-UV). The signal comes from $^{32}$P-end-labeling of RNA. Treatment of the precipitate with RNase I (+ RNase) reveals the size of the SR34a-GFP protein + ~5 kDa, corresponding to the short RNAs still bound by the protein after RNase treatment. The a-GFP immunoblot (bottom panel) identifies the precipitated SR34a-GFP protein.

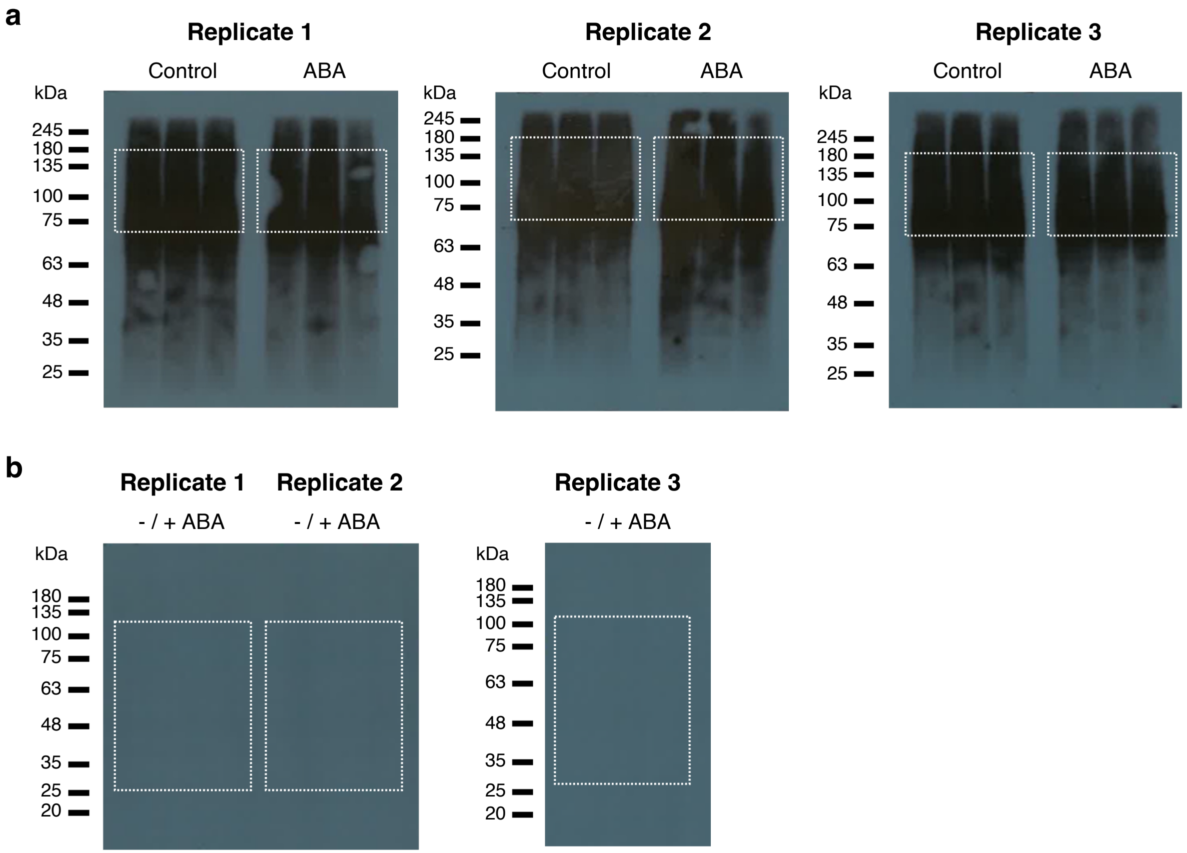


**Supplementary Fig. 6 | Autoradiograms of RNA-protein complexes isolated for iCLIP.** RNA-protein complexes immunoprecipitated from 44-h germinated seeds of the *SR34a:SR34a-GFP_sr34a-1* C1 (**a**) or *35S:GFP_sr34a-1* (**b**) transgenic lines treated or not with ABA. The rectangles indicate the regions that were cut out from the membrane for further RNA purification and sequencing.

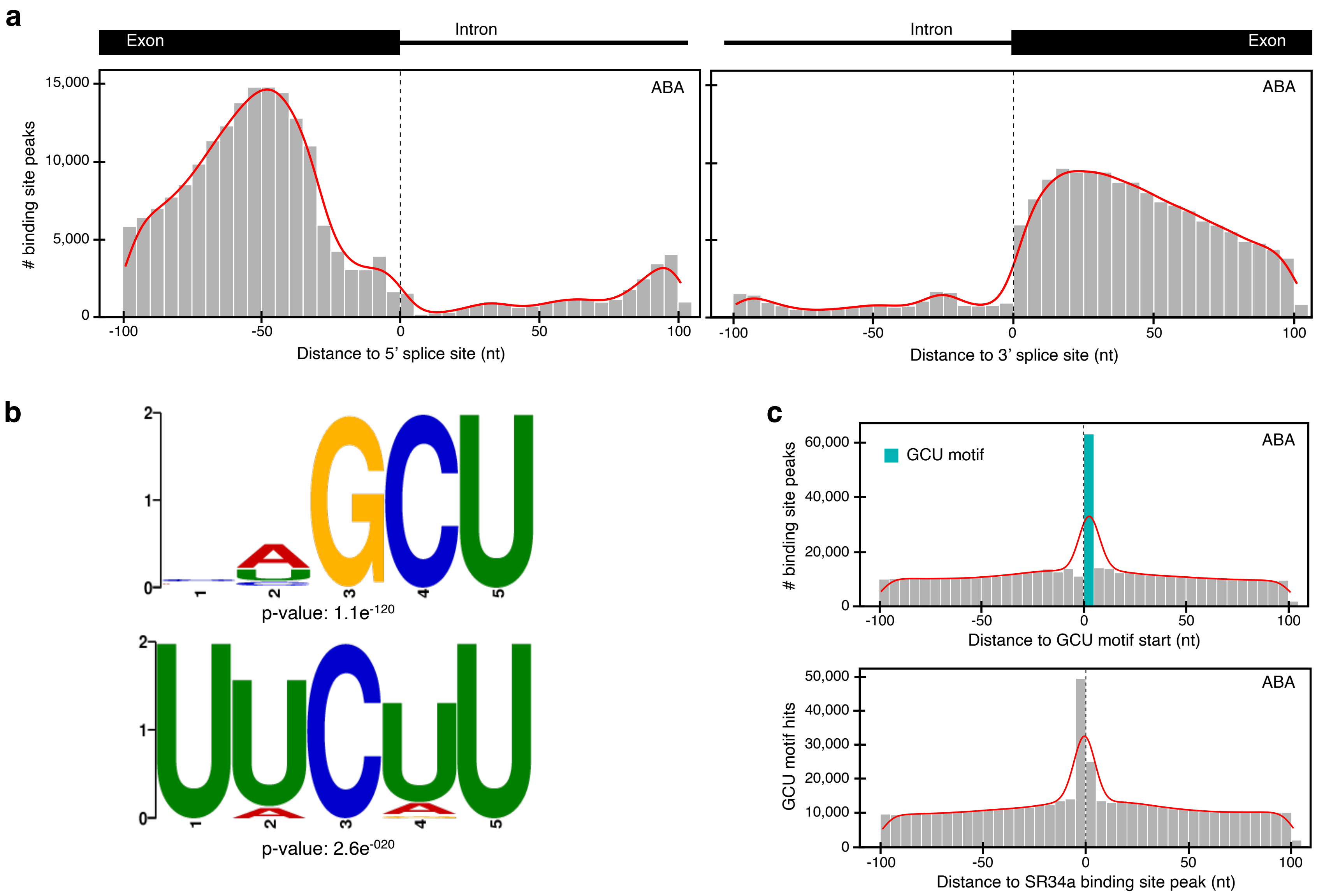


**Supplementary Fig. 7 | SR34a binding distribution and motifs under ABA conditions. a** SR34a iCLIP binding site peaks (i.e. middle position of binding sites) mapped in the vicinity of 5’ and 3’ splice sites under ABA conditions. **b** Binding motifs significantly enriched within SR34a binding sites identified under ABA conditions. **c** Number of iCLIP binding site peaks identified under ABA conditions as a function of their distance to GCU motifs in the *A. thaliana* genome (top), and number of GCU motif hits as a function of their distance to binding site peaks (bottom).

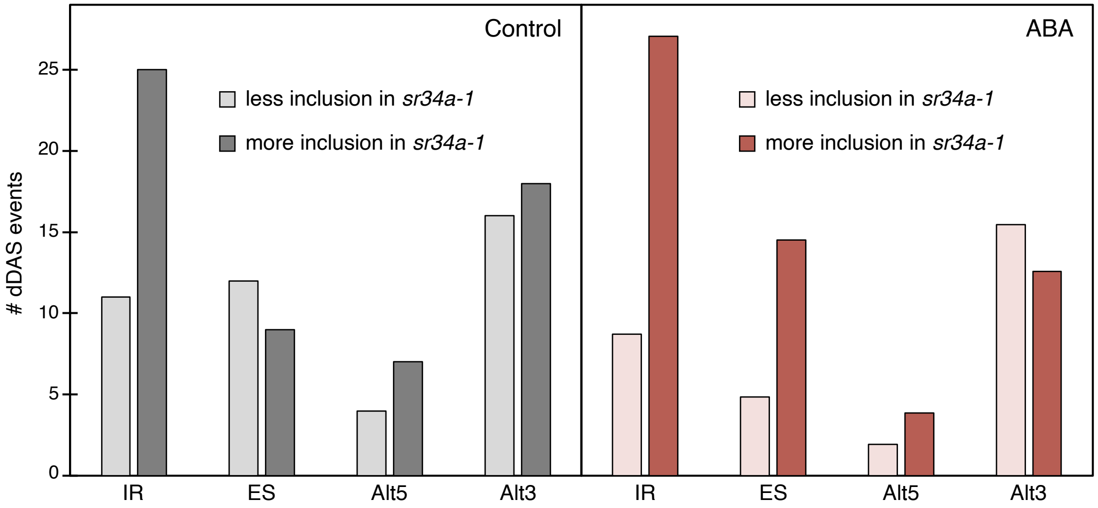


**Supplementary Fig. 8 | Distribution and regulation of the different types of alternative splicing event under direct SR34a control.** Number of intron retention (IR), exon skipping (ES), alternative 5' splice site (Alt5) and alternative 3' splice site (Alt3) events among the 102 or 92 dDAS events in the *sr34a-1* mutant identified under control (gray) or ABA (red) conditions, respectively. The lighter and darker colored bars indicate the number of dDAS events in which the alternative sequence was respectively less or more included in the *sr34a-1* mutant.
